# Lysosomal activity regulates *Caenorhabditis elegans* mitochondrial dynamics through vitamin B12 metabolism

**DOI:** 10.1101/2020.04.20.049502

**Authors:** Wei Wei, Gary Ruvkun

## Abstract

Mitochondrial fission and fusion are highly regulated by energy demand and physiological conditions to control the production, activity, and movement of these organelles. Mitochondria are arrayed in a periodic pattern in *Caenorhabditis elegans* muscle, but this pattern is disrupted by mutations in the mitochondrial fission component dynamin. Here we show that the dramatically disorganized mitochondria caused by a mitochondrial fission-defective dynamin mutation is strongly suppressed to a more periodic pattern by a second mutation in lysosomal biogenesis or acidification. Vitamin B12 is normally imported from the bacterial diet via lysosomal degradation of B12-binding proteins and transport of vitamin B12 to the mitochondrion and cytoplasm. We show that the lysosomal dysfunction induced by gene inactivations of lysosomal biogenesis or acidification factors causes vitamin B12 deficiency. Growth of the *C. elegans* dynamin mutant on an *E. coli* strain with low vitamin B12 also strongly suppressed the mitochondrial fission defect. Of the two *C. elegans* enzymes that require B12, gene inactivation of methionine synthase suppressed the mitochondrial fission defect of a dynamin mutation. We show that lysosomal dysfunction induced mitochondrial biogenesis which is mediated by vitamin B12 deficiency and methionine restriction. S-adenosylmethionine, the methyl donor of many methylation reactions, including histones, is synthesized from methionine by S-adenosylmethionine synthase; inactivation of the *sams-1* S-adenosylmethionine synthase also suppresses the *drp-1* fission defect, suggesting that vitamin B12 regulates mitochondrial biogenesis and then affects mitochondrial fission via chromatin pathways.

**SIGNIFICANCE STATEMENT:** The balance of mitochondrial fission and fusion, two aspects of mitochondrial dynamics, is important for mitochondrial function. Here we show that *Caenorhabditis elegans* lysosomal activity regulates mitochondrial dynamics by affecting mitochondrial fission through interfering the metabolism of a micronutrient, vitamin B12. Vitamin B12 is exclusively obtained from diets in animals including *C. elegans* and humans, and its uptake is mediated by the lysosome. We show that lysosomal dysfunction causes vitamin B12 deficiency that leads to reduction of methionine and S-adenosylmethionine to in turn increase mitochondrial biogenesis and fission. Our study provides an insight on the interactions between mitochondrial function and micronutrient metabolism.

## INTRODUCTION

The mitochondrion generates energy, supplies biosynthetic intermediates to carbohydrate, amino acid, steroid, and fat metabolic and anabolic pathways, and produces iron sulfur clusters. The various demands of metabolic intermediates and ATP in different cell types and under different conditions has led to a complex regulation of mitochondrial production, localization, and dynamics. Mitochondria are highly dynamic and undergo regulated fusion and fission to generate mitochondria in particular cells or subcellular locations where energy needs are high, or to scavenge mitochondria damaged by free radicals for example. Fusion can sustain mitochondrial function under aging and other conditions that damage mitochondria because fused mitochondria can compensate for organelle damage by sharing functional components (including mitochondrial DNA, RNAs, and proteins); fission contributes to quality control by segregating damaged mitochondria for degradation and also facilitating new mitochondria generation. The balance between fission and fusion may be disrupted during aging as age-dependent neurodegenerative diseases disrupt mitochondrial dynamics (1-3).

Mitochondrial fusion and fission are mediated by evolutionally conserved dynamin family guanosine triphosphatases (GTPases). *C. elegans* DRP-1 (DRP, dynamin-related protein) is a cytosolic dynamin that mediates mitochondrial fission, during which DRP-1 is recruited to the mitochondrial outer membrane to form spirals that constrict and sever the organelle (4-6). Two other *C. elegans* GTPases FZO-1 (Mitofusion 1, or MFN1 in mammals) and EAT-3 (Optic Atrophy 1, or OPA1 in mammals) mediate the opposite activity of the fission/fusion cycle of mitochondria, fusion of mitochondrial outer and inner membranes, respectively. Other receptors and mediators have been identified by genetic analysis in yeast of the pathways for mitochondrial fusion and fission, although many are not conserved in animals (1-3).

Here we show that a dynamin mutation that disrupts mitochondrial fission causes a dramatic tangling of the normally periodic pattern of mitochondria in *C. elegans* muscle cells, and that many gene inactivations or drugs that cause lysosomal dysfunction strongly suppress these mitochondrial defects to a highly periodic pattern. The lysosome is also a membrane-bound subcellular organelle. It serves as the scavenger of the cell to degrade components, and in defense against bacteria and viruses. As an intracellular digestive system, its function requires the acidification of the lysosomal lumen via a membrane-anchored proton pump, the vacuolar H^+^-ATPase (V-ATPase) (7) to in turn activate acid proteases. These acidified vesicles contain hydrolytic enzymes that can break down various macromolecules. The lysosome also delivers small molecule nutrients by degradation of extracellular or intracellular substrates. For example, in mammals vitamin B12 is retrieved from the diet by the lysosome (Nielsen et al., 2012). We find that in *C. elegans*, dietary vitamin B12 deficiency, similar to lysosomal dysfunction, can also strongly suppress the mitochondrial dynamics defect of a mitochondrial fission mutant. Dietary B12 supplementation effectively reduces the mitochondrial abnormalities, developmental and locomotion defects in lysosomal defective animals. Vitamin B12, or cobalamin, is an essential animal micronutrient that is exclusively synthesized by bacteria and archaea (8, 9). Within animal cells, B12 is a cofactor for methionine synthase (MTR) that catalyzes the methylation of homocysteine to methionine in the cytosol (10); and methylmalonyl-CoA mutase (MCM), that catalyzes the methylmalonyl-CoA to succinyl-CoA reaction in the catabolism of branched chain amino acids, odd-chain fatty acids, and cholesterol in the mitochondrion (11). These enzymes are encoded by *C. elegans metr-1* and *mmcm-1*, respectively. We found that inactivation of *metr-1* but not *mmcm-1* recapitulated the suppression of dynamin fission defects by B12 deficiency. Thus, vitamin B12 deficiency, either caused by lysosomal dysfunction or vitamin B12 dietary deficiency decreases the activity of methionine synthase, which leads to induce mitochondrial biogenesis that is coupled to an increase of fission activity for new mitochondrial generation, to suppress the loss of dynamin-mediated mitochondrial fission.

## RESULTS

### Inactivation of *spe-5* vacuolar ATPase suppresses the lethality and mitochondrial fission defects of a *drp-1/*mitochondrial dynamin mutant

*drp-1(or1393)* is a temperature-sensitive allele with a conserved glycine to glutamic acid (G39E) substitution mutation in the N-terminal dynamin domain. This mutation causes high penetrance lethality at the non-permissive temperature (12) (Figs. 1A, 1B and S1A). Because DRP-1 is a key mediator of mitochondrial fission, gene activities that regulate DRP-1 activity or the process of mitochondrial fission or fusion may affect the penetrance of *drp-1(or1393)* lethality. We performed an RNAi screen to identify gene inactivations that suppress the fission defects of the *drp-1(or1393)* mutant (Fig. S1B). We took advantage of the similarity between the mitochondrial fission defects of a *drp-1* mutant and a dynein heavy chain gene *dhc-1* mutant; gene inactivation of *dhc-1* causes mitochondrial fission defects that disrupt the periodic pattern of muscle mitochondria as strongly as a *drp-1* gene inactivation or mutation (Fig. S1C), suggesting that DHC-1 also functions in mitochondrial fission. A genome-scale RNAi screen for suppression of *dhc-1* lethality identified 49 gene inactivations (13) that were good candidates to also suppress *drp-1* lethality and mitochondrial fission defects. We tested the 49 *dhc-1* suppressor gene inactivations on *drp-1(or1393)* (Fig. S1B). Of these 49 candidate genes, we found that inactivations of *spe-5/*vacuolar ATPase and C42C1.3 partially suppressed the lethality of *drp-1(or1393)* at 23°C (Fig. S1D). Inactivation of *spe-5/*vacuolar ATPase increased ∼ 5 fold the viability of *drp-1(or1393)* at the non-permissive temperature 23°C, as well as at 15°C and 20°C, but did not affect wild-type (Figs. 1A and 1B). *spe-5* inactivation also suppressed the lethality of another widely used loss-of-function allele, *drp-1(tm1108)* (Fig. S1E). C42C1.3 is uncharacterized and worm-specific. We did not further study this hit.

**Fig. 1.**
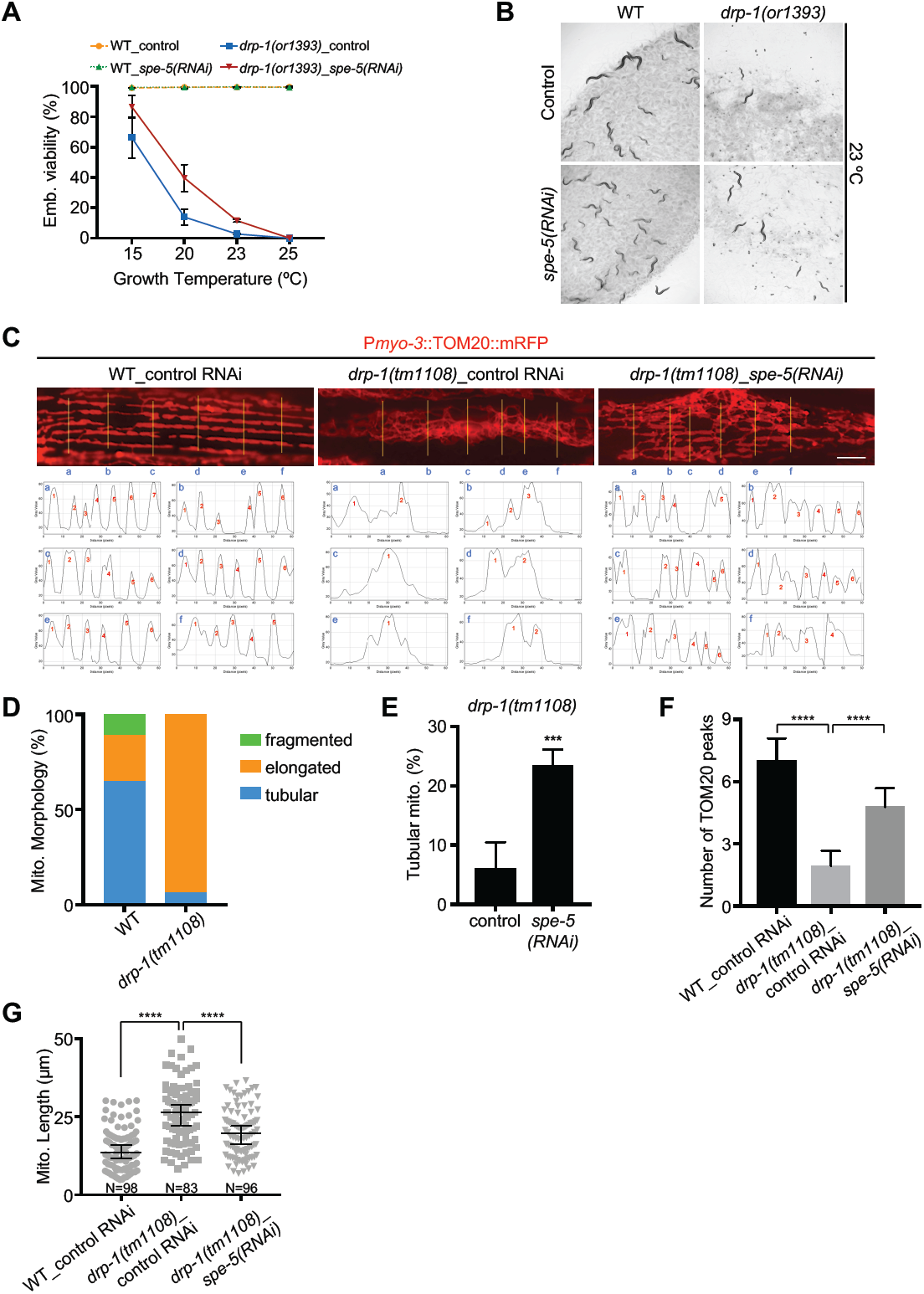
Inactivation of the vacuolar ATPase *spe-5* suppresses *drp-1* lethality and mitochondrial fission defects. (*A, B*) Viability of wild-type (WT) and *drp-1(or1393)* in control versus *spe-5(RNAi)* at indicated temperatures. Scale bar, 1 mm. (*C*) Mitochondrial morphology of wild-type and *drp-1(tm1108)* in control versus *spe-5(RNAi)* in a single body wall muscle cell. Mitochondria are visualized by a mitochondrial outer membrane localized mRFP fusion protein. Lower panels are plot profiles of the cross sections indicated by yellow lines. Criterion for a peak: peak to trough (both sides) > 20 Grey value units (y units). Scale bar, 5 μm. (*D)* Percentage of mitochondria with fragmented, elongated or tubular morphology in wild-type and *drp-1(tm1108)* in body wall muscles. (*E)* Percentage of mitochondria with tubular morphology in *drp-1(tm1108)* body wall muscles after indicated RNAi treatments. Mean ± s.d. \*\*\**P* < 0.001. (*F)* Peak numbers for the plot profiles of mitochondrial morphology in indicated animals. Mean ± s.d. \*\*\*\**P* < 0.0001. N=40∼50 for each group. (*G)* Mitochondrial lengths in body wall muscles in indicated animals. Mitochondrial lengths were calculated by MiNA toolset. Median with 95% C. I. Mann-Whitney test. \*\*\*\**P* < 0.0001. N indicates the sample size.

Mitochondria in any cell of body muscle of wild type animals can be visualized with mitochondrial RFP fusion proteins that form a highly periodic two-dimensional network, with strong fluorescent peaks ∼ 0.4 microns wide and ∼ 13.5 microns long, where mitochondria are localized along sarcomere, and troughs ∼ 1.3 microns wide with no fluorescence that have no mitochondria (Fig. 1C). In wild type, muscle mitochondria are localized in multiple parallel arrays along the sarcomeres of muscles, as visualized in one muscle cell in each panel of Figure 1C, for example, by an RFP protein fused at the N-terminus of the translocase of outer mitochondrial membrane TOMM-20 (Figs. 1C and 1D). The *drp-1* mutation causes defects in mitochondrial fission so that mitochondria fuse more than normal into an elongated and very tangled, non-parallel mitochondrial morphology in more than 90% of muscle mitochondria (Figs. 1C and 1D). Inactivation of *spe-5* strongly suppressed the mitochondrial fission defects in *drp-1(tm1108)*; ∼ 25% of mitochondria were restored to a highly parallel, tubular-like morphology that are more like wild-type (Figs. 1C, 1E and S2). To more precisely quantitate a periodic or aperiodic mitochondrial network in suppressed mitochondrial fission mutants or the mitochondrial fission mutant, respectively, we profiled the pattern of fluorescence across six cross sections per muscle cell image, to display the periodic placement of mitochondria. While highly parallel, tubular-like mitochondria of wild type *C. elegans* displayed periodic TOMM-20 peaks from the plot profiles of the cross sections of the morphology images, the elongated and very tangled mitochondria of *drp-1* fission defective animals showed much fewer TOMM-20 peaks from the plot profiles of mitochondrial morphology images (Figs. 1C and S2). We found that *drp-1(tm1108)* animals had ∼ 1.9 TOMM-20 peaks on average in each muscle cell image, much fewer than wild-type animals which showed ∼ 7.0 TOMM-20 peaks on average. Inactivation of *spe-5* increased the average TOMM-20 peaks to almost 5 peaks per cells in *drp-1(tm1108)*, or about 2.5x more than *drp-1(tm1108)* (Fig. 1F), suggesting the mitochondrial fission defect in *drp-1(tm1108)* was ameliorated by *spe-5* inactivation. An independent measure of the suppression of fission defects was to determine mitochondrial lengths. The mitochondria are highly tangled in *drp-1(tm1108)*, which makes it impossible to measure the mitochondrial lengths from simple photographic images. We used a Mitochondrial Network Analysis (MiNA) toolset to generate a morphological skeleton for the mitochondrion to calculate the mitochondrial lengths by ImageJ (14). We found that increased mitochondrial lengths by DRP-1 dysfunction was suppressed by *spe-5* inactivation (Fig. 1G). These results suggest the involvement of *spe-5/*vacuolar ATPase in mitochondrial fission regulation.

Mitochondrial function is tightly surveilled in eukaryotes. If mitochondrial dysfunction is detected, cells activate a series of responses including the mitochondrial unfolded protein response (mtUPR). HSP-6 is one of the mitochondrial chaperones that mediates the mtUPR (15, 16). P*hsp-6*::GFP was strongly induced in *drp-1(tm1108)* (Fig. S3A). If *spe-5* inactivation mitigated the mitochondrial fission defects of *drp-1*, it should also alleviate the mtUPR induced by *drp-1* deficiency. Indeed, gene inactivation of *spe-5* reduced P*hsp-6*::GFP induction two fold in *drp-1(tm1108)* (Fig. S3A). In contrast, *spe-5* depletion did not decrease and in fact slightly increased P*hsp-6*::GFP expression in wild-type (Fig. S3A). Consistent with the reporter gene, the induction of endogenous mitochondrial chaperone genes *hsp-6* and *hsp-60* by *drp-1* depletion were suppressed by *spe-5* inactivation. (Fig. S3B). Consistent with the lower mitochondrial stress in the *drp-1(tm1108); spe-5 (RNAi)* strain, we found that inactivation of *spe-5* increased mitochondrial membrane potential in *drp-1(tm1108)* (Fig. S3C).

### V-ATPase dysfunction suppresses *drp-1* lethality and mitochondrial fission defects

*spe-5* encodes a subunit of the vacuolar V-ATPase (17), suggesting that V-ATPase activity or the activity of the lysosome might regulate mitochondrial dynamics. To test if lysosomal function was needed, we depleted other V-ATPase subunits and found that inactivations of other subunit genes also suppressed *drp-1* lethality (Fig. 2A). Consistent with the lethality suppression, these V-ATPase gene inactivations restored the tangled non-periodically aligned mitochondria of the *drp-1(tm1108)* mutant to a more wild type parallel, tubular-like morphology (Figs. 2B and 2C). The suppression by V-ATPase gene inactivations was most obvious in the number of TOMM-20 peaks from the plot profiles of mitochondrial morphologies which increased ∼ 3 fold by V-ATPase gene inactivations in *drp-1(tm1108)* (Fig. 2D). Consistent with this periodicity analysis, V-ATPase gene inactivations decreased mitochondrial lengths in *drp-1(tm1108)* (Fig. 2E). Drug inhibition of vacuolar V-ATPase activity with bafilomycin A1 (BafA1) (18) or concanamycin A (CMA) (19) also suppressed *drp-1* lethality but had no effect on wild-type (Figs. S4A and S4B). BafA1 and CMA treatment suppressed the mitochondrial fission defects in *drp-1(tm1108)*, increasing the percentage of mitochondria with a parallel tubular morphology by ∼ 2 fold (Figs. 2F and 2G). Similarly, BafA1 and CMA treatment increased the number and periodicity of TOMM-20 peaks from the plot profiles of mitochondrial morphologies (Fig. 2H) and decreased mitochondrial lengths in *drp-1(tm1108)* (Fig. 2I), suggesting mitochondrial fission defects in *drp-1(tm1108)* were mitigated by these V-ATPase activity inhibitors. Disrupting V-ATPase activity by gene inactivation or chemical inhibitor treatment decreased the induction of P*hsp-6*::GFP and endogenous mitochondrial chaperone genes *hsp-6* and *hsp-*60 in *drp-1(tm1108)* (Figs. S4C-S4F), suggesting that the mitochondrial defects normally caused by a decrease in *drp-1* activity are ameliorated by also inducing vacuolar V-ATPase dysfunction. V-ATPase dysfunction by gene inactivation or chemical inhibitor treatment increased mitochondrial membrane potential in *drp-1(tm1108)* (Figs. S5A and S5B), suggesting that mitochondrial functions were improved. Overall, these data show that the activity of the key proton pump of the lysosome, the vacuolar V-ATPase, regulates the mitochondrial over-fusion defect caused by DRP-1 dynamin-mediated mitochondrial fission dysfunction.

**Fig. 2.**
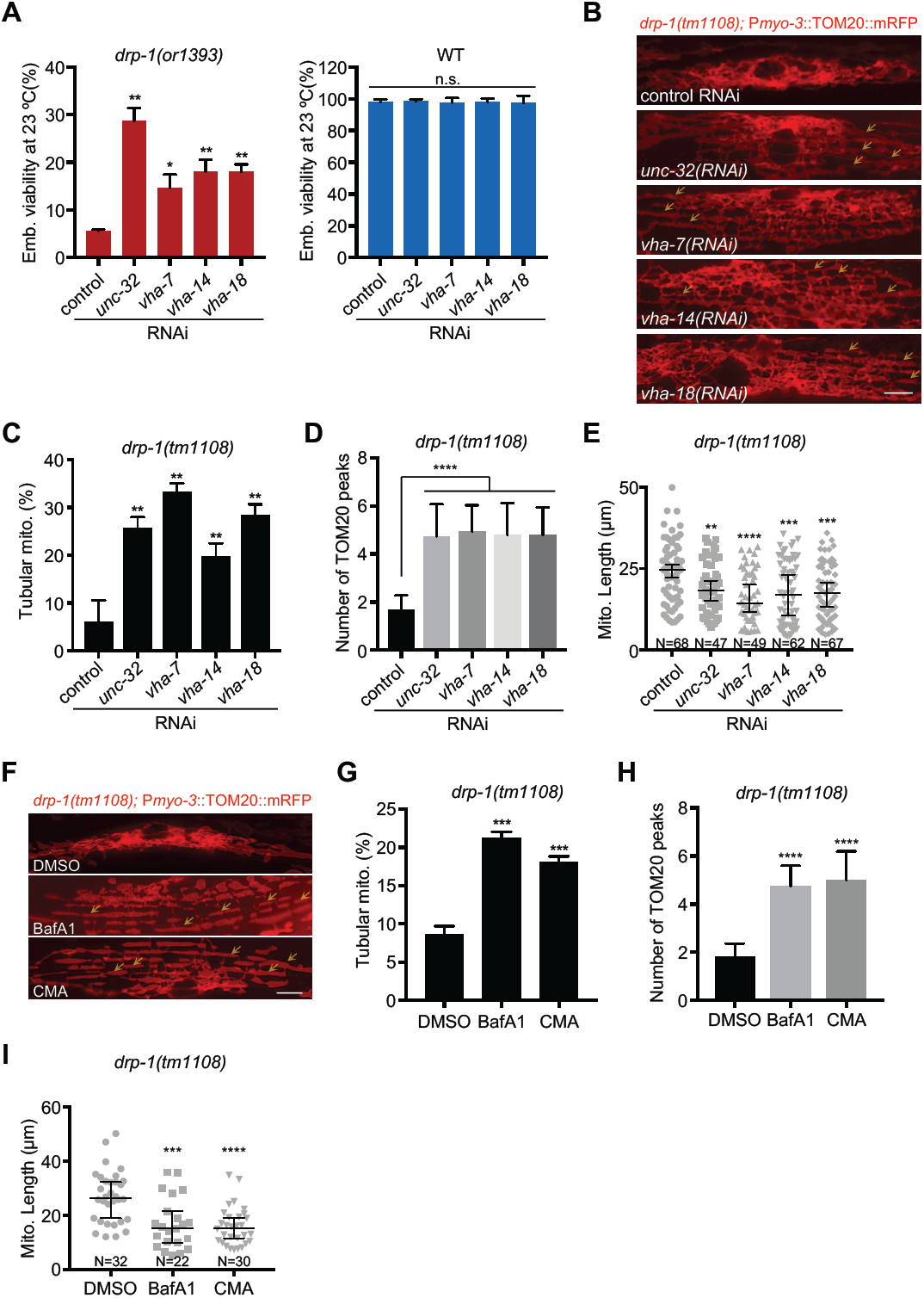
V-ATPase dysfunction suppresses *drp-1* lethality and mitochondrial fission defects. (*A)* Viability of wild-type (WT) or *drp-1(or1393)* after V-ATPase gene inactivations at 23°C. Mean ± s.d. \*\**P* < 0.01, \**P* < 0.05. (*B)* Mitochondrial morphology in *drp-1(tm1108)* body wall muscles after indicated RNAi treatment. Arrows indicate some examples of nice parallel mitochondrial structure. Scale bar, 5 μm. (*C)* Percentage of mitochondria with tubular morphology in *drp-1(tm1108)* body wall muscles after indicated RNAi treatments. Mean ± s.d. \*\**P* < 0.01. (*D)* Peak numbers for the plot profiles of mitochondrial morphology in indicated animals. Mean ± s.d. \*\*\*\**P* < 0.0001. N=40∼50 for each group. (*E)* Mitochondrial lengths in body wall muscles in indicated animals. Mitochondrial lengths were calculated by MiNA toolset. Median with 95% C. I. Mann-Whitney test. \*\*\*\**P* < 0.0001, \*\*\**P* < 0.001, \*\**P* < 0.01. N indicates the sample size. (*F)* Mitochondrial morphology in *drp-1(tm1108)* body wall muscles with DMSO, BafA1 or CMA treatment. DMSO, solvent control; BafA1, bafilomycin A1; CMA, concanamycin A. Arrows indicate some examples of nice parallel mitochondrial structure. Scale bar, 5 μm. (*G)* Percentage of mitochondria with tubular morphology in *drp-1(tm1108)* body wall muscles after DMSO, BafA1 or CMA treatment. Mean ± s.d. \*\*\**P* < 0.001. (*H)* Peak numbers for the plot profiles of mitochondrial morphology in *drp-1(tm1108)* body wall muscles after DMSO, BafA1 or CMA treatment. Mean ± s.d. \*\*\*\**P* < 0.0001. N=40∼50 for each group. (*I)* Mitochondrial lengths in body wall muscles in *drp-1(tm1108)* body wall muscles after DMSO, BafA1 or CMA treatment. Mitochondrial lengths were calculated by MiNA toolset. Median with 95% C. I. Mann-Whitney test. \*\*\*\**P* < 0.0001, \*\*\**P* < 0.001. N indicates the sample size.

### Lysosomal activity affects mitochondrial dynamics

V-ATPase is the proton pump that acidifies lysosome. To further test the involvement of the lysosome in mitochondrial dynamics, we examined more factors that regulate lysosomal biogenesis or activity. HLH-30, the *C. elegans* orthologue of the transcription factor EB (TFEB), is a master regulator of lysosomal biogenesis (20). LMP-1 (LAMP-1 in mammals) and LMP-2 (LAMP-2 in mammals) (<u>L</u>ysosomal-<u>a</u>ssociated <u>M</u>embrane <u>P</u>rotein 1 and 2, respectively) constitute ∼ 50% of the lysosomal membrane proteins and are required for lysosomal biogenesis and maintaining lysosome integrity (21, 22). We found that inactivations of *hlh-30, lmp-1*, and *lmp-2* each suppressed the mitochondrial fission defects caused by *drp-1* deficiency, increasing the percent of mitochondria with a parallel tubular morphology by 4-5 fold (Figs. 3A and 3B). The numbers of TOMM-20 peaks from the plot profiles of mitochondrial morphologies in *drp-1(tm1108)* were increased by ∼ 3 fold by inactivations of *hlh-30, lmp-1*, or *lmp-*2 (Fig. 3C). Consistent with the periodic mitochondrial morphology analysis, these lysosomal gene inactivations decreased mitochondrial lengths in *drp-1(tm1108)* (Fig. 3D), suggesting that mitochondrial fission defect in *drp-1(tm1108)* was ameliorated by lysosomal dysfunction. Inactivations of *hlh-30, lmp-1* and *lmp-2* mildly suppressed the induction of P*hsp-6*::GFP and endogenous *hsp-6* and *hsp-*60 expression in *drp-1(tm1108)* (Figs. S6A and S6B), suggesting that the mitochondrial homeostasis in *drp-1(tm1108)* were improved by disruption of lysosomal activity. Similarly, lysosomal dysfunction by inactivations of *hlh-30, lmp-1*, or *lmp-*2 increased mitochondrial membrane potential in *drp-1(tm1108)* (Fig. S6C), suggesting improvement of mitochondrial function.

**Fig. 3.**
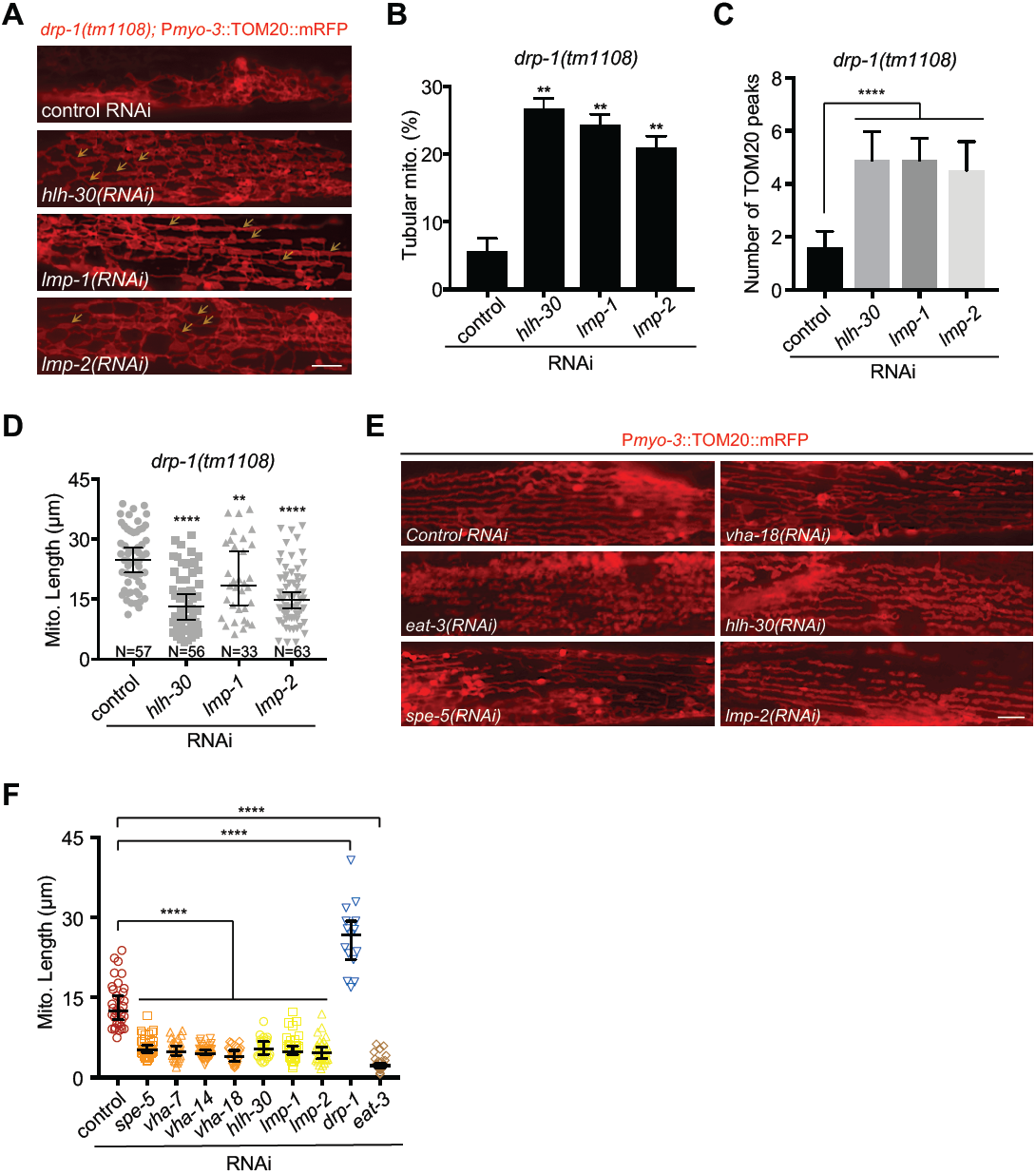
Lysosomal activity affects mitochondrial dynamics. (*A)* Mitochondrial morphology in *drp-1(tm1108)* body wall muscles after indicated RNAi treatment. Arrows indicate some examples of nice parallel mitochondrial structure. Scale bar, 5 μm. (*B)* Percentage of mitochondria with tubular morphology in *drp-1(tm1108)* body wall muscles after indicated RNAi treatments. Mean ± s.d. \*\**P* < 0.01. (*C)* Peak numbers for the plot profiles of mitochondrial morphology in indicated animals. Mean ± s.d. \*\*\*\**P* < 0.0001. N=40∼50 for each group. (*D)* Mitochondrial lengths in body wall muscles in indicated animals. Mitochondrial lengths were calculated by MiNA toolset. Median with 95% C. I. Mann-Whitney test. \*\*\*\**P* < 0.0001, \*\**P* < 0.01. N indicates the sample size. (*E)* Mitochondrial morphology in body wall muscles after indicated RNAi treatment. Scale bar, 5 μm. (*F)* Lengths of mitochondria after indicated gene inactivations by RNAi. n=15∼40, Median with 95% C. I. Mann-Whitney test. \*\*\*\**P* < 0.0001.

We speculated that lysosomal activity might act in mitochondrial fission, for example, by lysosomal-mediated mitophagy. However, we found that inactivations of the mitophagy pathway genes including the mitochondrial phosphatase and tensin (PTEN)-induced kinase 1 (PINK-1), the cytosolic E3 ubiquitin ligase Parkin (PDR-1) or the Atg6/Vps30/Beclin1 homologue BEC-1 did not suppress the lethality or mitochondrial fission defects in *drp-1* defective animals (Figs. S6D-S6F). It makes sense that mitophagy defect does not affect mitochondrial morphology in *drp-1* depleted animals because DRP-1-dependent mitochondrial fission is a prerequisite for mitophagy (23) so that the dysfunctions of the downstream mitophagy could not rescue the upstream DRP-1 defect in mitochondrial morphology.

Because either inhibition of vacuolar V-ATPase activity or disruption of lysosomal biogenesis/integrity suppresses the mitochondrial fission defects of *drp-1/*dynamin mutant, a reasonable model is that lysosomal dysfunction promotes mitochondrial fission so as to re-balance the fusion-fission events in the *drp-1* mutant. Indeed, we found that inactivations of V-ATPase subunit genes or lysosomal biogenesis/integrity genes caused a modest increase in mitochondrial fragmentation in an otherwise wild type background (Figs. 3E and 3F). However, the mitochondrial fragmentation caused by lysosomal dysfunction was less severe than that caused by disruption of *eat-3* that encodes the homologue of the mitochondrial inner membrane fusion dynamin OPA1 (Figs. 3E and 3F), suggesting that lysosome does not directly function in mitochondrial fission/fusion, and/or other regulatory pathways are also involved.

### Vitamin B12 deficiency increases mitochondrial fission and mitigates *drp-1* defects

*E. coli* bacteria are the standard laboratory food source for *C. elegans*. The most common *C. elegans* diet in lab is OP50, an *E. coli* B-type strain (24), whereas an *E. coli* K12-derived RNase III deficient strain HT115 is used for RNA interference screens. Most *C. elegans* genetic analysis and day to day maintenance of strains is done on OP50 *E. coli* B which is then switched to HT115 for feeding RNAi experiments. We noticed that the *drp-1* loss-of-function (*lf*) mutant, *drp-1(tm1108)* (*drp-1(lf)* hereafter) grew faster and with more viability on *E. coli* OP50 than on *E. coli* HT115 (Figs. 4A and S7A). *drp-1(lf)* grown on *E. coli* OP50 induced P*hsp-6*::GFP less than those grown on *E. coli* HT115 (Fig. S7B, upper panels), suggesting that *E. coli* B supplies or lacks a key molecule that mediates the mitochondrial abnormalities in *drp-1(lf)*. Consistent with the improved growth and viability on *E. coli* OP50, the highly tangled mitochondria in *drp-1(lf)* was suppressed by feeding on *E. coli* OP50 compared to *E. coli* HT115 (Figs. 4B, upper panels, and 4C). The TOMM-20 peaks from the plot profiles of mitochondrial morphologies in *drp-1(lf)* were increased by 2.4 fold by feeding on *E. coli* OP50 compared to *E. coli* HT115 (Fig. 4D), suggesting that mitochondrial fission defect in *drp-1(lf)* was ameliorated by feeding on *E. coli* OP50 diet compared to *E. coli* HT115.

**Fig. 4.**
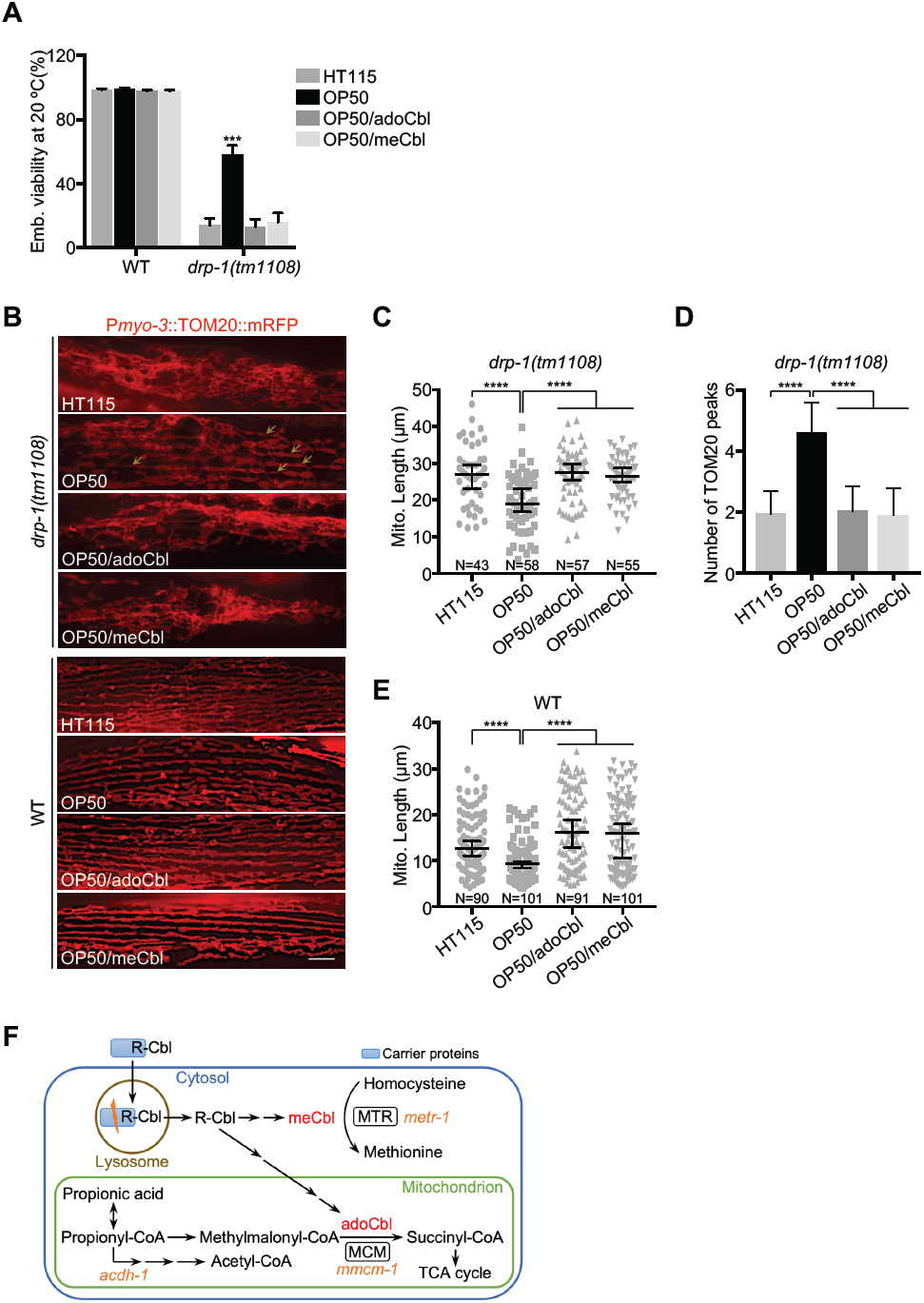
Vitamin B12 deficiency increases mitochondrial fission and mitigates *drp-1* defects. (*A)* Viability of wild-type (WT) or *drp-1(tm1108)* fed on *E. coli* HT115 (B12 proficient) or *E. coli* OP50 (B12 deficient) with or without B12 supplementation. adoCbl, adenosylcobalamin; meCbl, methylcobalamin. The final concentration of adoCbl or meCbl supplemented to the culture plates is 12.8 nM. Mean ± s.d. \*\*\**P* < 0.001. (*B)* Mitochondrial morphology in body wall muscles in *drp-1(tm1108)* (upper panels) or wild-type (lower panels) fed on *E. coli* HT115 or *E. coli* OP50 with or without B12 supplementation. Arrows indicate some examples of nice parallel mitochondrial structure in *drp-1(tm1108)* on *E. coli* OP50. Scale bar, 5 μm. (*C, E*) Mitochondrial lengths in body wall muscles in *drp-1(tm1108)* (D) or wild-type (E) animals fed on *E. coli* HT115 or *E. coli* OP50 with or without B12 supplementation. Mitochondrial lengths were calculated by MiNA toolset. Median with 95% C. I. Mann-Whitney test. \*\*\*\**P* < 0.0001. N indicates the sample size. (*D*) Peak numbers for the plot profiles of mitochondrial morphology in *drp-1(tm1108)* animals fed on *E. coli* HT115 or *E. coli* OP50 with or without B12 supplementation. Mean ± s.d. \*\*\*\**P* < 0.0001. N=40∼50 for each group. (*F*) Vitamin B12 metabolic pathways. MTR, methionine synthase; MCM, methylmalonyl-CoA mutase. *C. elegans* genes are in orange italics.

There are many genetic differences between *E. coli* B and *E. coli* K12, but one difference that has been noted is that *E. coli* B supplies low amounts of vitamin B12 to *C. elegans* (25-27). *E. coli* strains do not carry the biosynthetic operons for B12 synthesis but hundreds of disparate bacterial species synthesize vitamin B12 (28). In many animals, including *C. elegans* and humans, vitamin B12 is exclusively obtained from bacteria or diets and is necessary for a variety of methyl transfer reactions. *E. coli* strains in natural ecosystems obtain vitamin B12 from other organisms. In the lab, *E. coli* transports vitamin B12 from the LB growth media. However, the *E. coli* B-type strain has greatly reduced vitamin B12 transport and uptake from the growth medium, which is usually supplemented with peptone to provide vitamins and other nutrients essential for growth (29-31). *E. coli* K12 appears to take up higher levels of vitamin B12 from the bactopeptone growth media on worm plates to supply to the animals (25-27).

As a micronutrient, the endogenous vitamin B12 level is very low and thus hard to quantify directly and accurately. However, B12 deficiency compromises the methylmalonyl-CoA mutase-mediated mitochondrial propionic acid breakdown pathway and activates a propionate shunt; the first step of this pathway is carried out by acyl-coenzyme A dehydrogenase ACDH-1 (Fig. 4F) (Watson et al., 2016). P*acdh-1*::GFP has been used as a reporter of cellular B12 supply (26, 27). *C. elegans* grown on *E. coli* OP50 strongly induced P*acdh-1*::GFP, but this gene fusion is much more weakly expressed in animals feeding on *E. coli* HT115, a K12 strain (Figs. S8A, upper panels and S8C). This demonstrates the dietary B12 deficiency for animals on *E. coli* OP50 but not on *E. coli* HT115.

To test whether the discrepancies of *drp-1(lf)* on the *E coli* K12 HT115 (B12 proficient) and *E. coli* B OP50 (B12 deficient) diets is caused by the B12 supply but other differences between these two *E. coli* strains, we supplemented the *E. coli* OP50 diet with pure vitamin B12 adoCbl or meCbl, which are actually interconvertible within animal cells (27, 32, 33). We found that either form of B12 supplementation effectively complemented the dietary B12 deficiency on *E. coli* OP50, as indicated by P*acdh-1*::GFP expression (Figs. S8A, upper panels, S8C and S8D). Feeding on *E coli* HT115 (B12 proficient) diet or on *E. coli* OP50 supplemented by B12 also suppressed the P*acdh-1*::GFP induction, suggesting that possible differences in metabolizing B12 in these *E. coli* strains are not the cause of the different phenotypes we observed above. We found that either adoCbl or meCbl supplementation abolished the suppression of *drp-1(lf)* lethality by an *E. coli* OP50 diet (Fig. 4A). In addition, B12 supplementation suppressed the decrease of P*hsp-6*::GFP induction in *drp-1(lf)* grown on *E. coli* OP50 (Fig. S7B, upper panels). Consistent with suppression of mitochondrial UPRs, we observed that the suppression of the *drp-1(lf)* mitochondrial disorganization and fission defect by feeding on *E. coli* OP50 was abrogated by B12 supplementation (Figs. 4B-D), indicating that the *E. coli* OP50 diet-caused B12 deficiency leads to the better growth and mitochondrial improvement in *drp-1(lf)*.

We tested the effects of B12 deficiency on mitochondrial dynamics. We found that wild type animals grown on *E. coli* OP50 had mildly increased mitochondrial fragmentation, which could be complemented by exogenous B12 supplementation (Fig. 4B, lower panels), as has been observed (Revtovich et al., 2019). Moreover, P*hsp-6*::GFP was slightly induced on an *E. coli* OP50 diet, indicating mild mitochondrial dysfunction caused by B12 deficiency (Fig. S7B, lower panels). In addition, synchronized animals grown on *E. coli* OP50 were shorter than those on B12 plus diets (Fig. S7B, lower panels), suggesting development is mildly compromised under B12 deficiency, as noted previously (25, 27).

### Lysosomal retrieval of vitamin B12

In mammals, dietary vitamin B12 is first internalized by carrier proteins including intrinsic factor and transcobalamin, and then transported into the lysosome, where the carrier proteins are degraded by acid proteases to liberate B12 into the cytoplasm where it is modified to methylcobalamin or adenosylcobalamin (Fig. 4F) (34). The enzymes that use vitamin B12 as a cofactor are broadly conserved, methionine synthase (MTR) and methylmalonyl-CoA mutase (MCM) (Fig. 4F).

Lysosomal function is required for B12 supply in *C. elegans*. Treating animals with the V-ATPase inhibitors CMA or BafA1 induced P*acdh-1*::GFP on an *E. coli* HT115 B12 proficient diet (Fig. S9A). Lysosomal dysfunction by genetic mutations or inactivations of V-ATPase subunit genes or lysosomal biogenesis/integrity genes by RNAi (on *E. coli* HT115) also induced P*acdh-1*::GFP (Figs. S9B and S9C). The P*acdh-1*::GFP induction by V-ATPase inhibitors or lysosomal dysfunction on *E. coli* HT115 suggested cellular B12 deficiency, probably due to disruption of vitamin B12 release from lysosome. The *lmp-1(lf)* mutation caused developmental delay and induction of P*hsp-6*::GFP on an *E. coli* OP50 B12 deficient diet (Fig. S10A) but vitamin B12 supplementation rescued the developmental defect and *hsp-6* induction on an *E. coli* OP50 diet (Fig. S10A). The *lmp-1(lf)* lysosomal defective animals grown on *E. coli* OP50 had more fragmented mitochondria and decreased average mitochondrial lengths in body wall muscles than those grown on *E. coli* HT115; dietary supplementation of B12 increased the mitochondrial connectivity and reduced the fragmentation defect in *lmp-1(lf)* grown on *E. coli* OP50 (Fig. S10B).

In humans, lysosomal storage diseases caused by mutations in lysosomal proteins are associated with fragmented mitochondria and manifest as neurodegeneration. We found that *lmp-1(lf)* lysosomal defective animals showed locomotion defects including decreased movement velocity and increased movement tortuosity especially on the B12 deficient *E. coli* OP50 diet (Figs. S10C and S10D). Dietary B12 supplementation greatly reduced the locomotion defects in *lmp-1(lf)* (Figs. S10C and S10D), consistent with the observation of improved mitochondrial health in *lmp-1(lf)* with dietary B12 supplementation (Figs. S10A and S10B).

### Methionine restriction mediates B12 deficiency-caused mitochondrial fragmentation

Vitamin B12 is a cofactor for cytosolic methionine synthase (MTR) and mitochondrial methylmalonyl-CoA mutase (MCM) that are encoded by *C. elegans metr-1* and *mmcm-1*, respectively (Fig. 4F). We examined which enzyme mediates the B12 deficiency-caused mitochondrial fragmentation. Inactivation of *metr-1* but not *mmcm-1* completely recapitulated the mitochondrial phenotypes caused by B12 deficiency. In particular, inactivation of *metr-1* suppressed the lethality and slow growth in *drp-1(lf)* but delayed development in *lmp-1(lf)*, whereas RNAi of *mmcm-1* did not affect growth or mitochondrial morphology of *drp-1(lf)* or *lmp-1(lf)* (Figs. 5A, S11A and S11B, upper panels). Inactivation of *metr-1* in a wild-type background caused modest mitochondrial fragmentation, similar to lysosomal dysfunction or dietary B12 deficiency, while RNAi of *metr-1* reduced the mitochondrial fission defect in *drp-1(lf)* but enhanced the mitochondrial fragmentation defect in *lmp-1(lf)* fed on *E. coli* HT115 RNAi bacteria (Fig. 5B). Consistent with the mitochondrial status, P*hsp-6*::GFP induction is decreased in *drp-1(lf)* but increased in *lmp-1(lf)* by *metr-1* gene inactivation, while RNAi of *metr-1* mildly induced P*hsp-6*::GFP in an otherwise wild-type background (Fig. S11B, lower panels). Gene inactivation of *mmcm-1* did not affect mitochondrial morphology or *hsp-6* expression more than control RNAi (Figs. 5B and S11B, lower panels). Overall, these data indicate that B12 deficiency-caused mitochondrial fragmentation is mediated by the vitamin B12 -requiring enzyme methionine synthase.

**Fig. 5.**
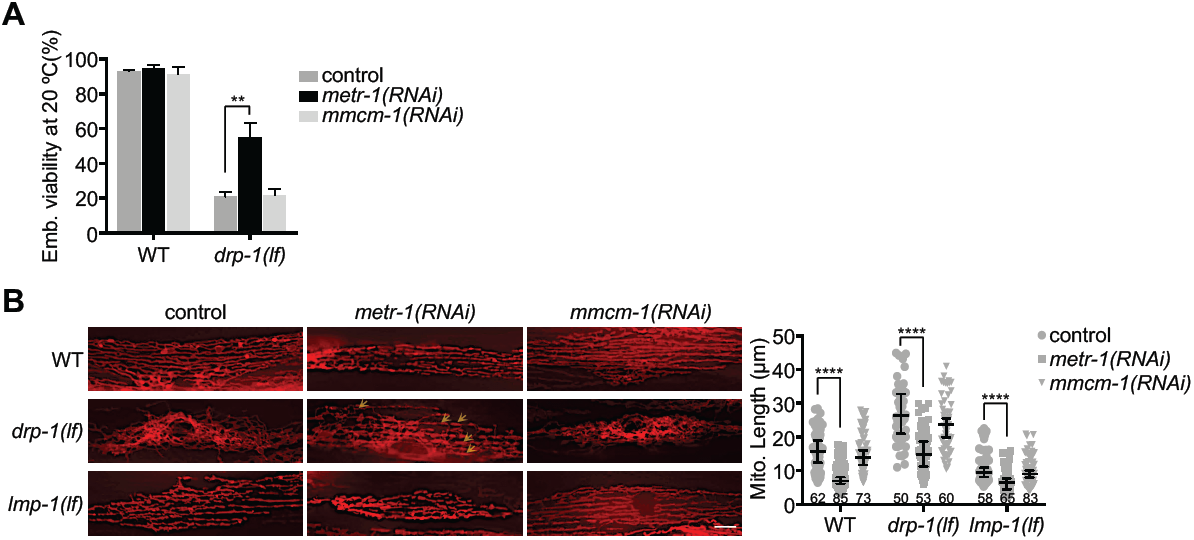
Inactivation of methionine synthase increases mitochondrial fission. (*A)* Viability of wild-type (WT) and *drp-1(lf)* grown on indicated RNAi clones. Mean ± s.d. \*\**P* < 0.01. (*B)* Mitochondrial morphologies and lengths in body wall muscles in wild-type, *lmp-1(lf)* and *drp-1(lf)* grown on indicated RNAi clones. Mitochondrial lengths were calculated by MiNA toolset. Median with 95% C. I. Mann-Whitney test. \*\*\*\**P* < 0.0001. N indicates the sample size. Scale bar, 5 μm.

Methionine synthase converts homocysteine to methionine that is required for the synthesis of methyl donor S-adenosylmethionine (SAM). Disruption of MTR enzymatic activity reduces methionine biosynthesis and leads to methionine restriction. We reasoned that the reduction of methionine mediated the increased mitochondrial fragmentation observed in lysosomal defective or B12 deficient animals. Indeed, dietary supplementation of methionine greatly reduced the P*hsp-6*::GFP induction by *metr-1* gene inactivation or dietary B12 deficiency in *lmp-1(lf)* (Fig. S11C, upper panels). Methionine supplementation also reduced the mild *hsp-6* induction by *metr-1* RNAi or dietary B12 deficiency in wild type animals (Fig. S11C, lower panels). Methionine restriction leads to decrease of SAM levels. To test if low SAM causes the mitochondrial phenotypes observed in B12 deficient or lysosomal deficient animals, we inactivated *sams-1* that encodes the S-adenosylmethionine synthetase and found that SAM reduction increased mitochondrial fission in wild-type as well as mitigated the mitochondrial fission defects in *drp-1(lf)* animals (Fig. S12A), similar to that caused by B12 deficiency or lysosomal dysfunction. Inactivation of *sams-1* mildly induced P*hsp-6*::GFP similar to *metr-1* RNAi (Fig. S12B). Mitochondrial fragmentation caused by *sams-1* inactivation could not rescued by exogenous B12 supplementation or methionine supplementation (Fig. S12C). Combined, these results indicate that methionine restriction by B12 deficiency leads to reduction of SAM, which causes an increase of mitochondrial fission.

### Lysosomal dysfunction augments mitochondrial biogenesis

Methionine restriction increased mitochondrial DNA (mtDNA) copy number and mitochondrial density in some rat tissues (35, 36). We found that deletion of *C. elegans metr-*1 increased mtDNA level 1.5 fold compared to wild-type. Dietary methionine supplementation reduced mtDNA levels either in wild-type or *metr-1(lf)* (Fig. 6A). Nonyl Acridine Orange (NAO) is a fluorescent marker that is retained in mitochondria independent of the mitochondrial membrane potential, making it an ideal probe to measure mitochondrial mass (37). Mitochondrial mass measured by NAO is comparable between wild-type and *metr-1(lf)*. The discrepancy might be due to differential growth rates, as *metr-1(lf)* showed mild developmental delay compared to wild-type. However, methionine supplementation significantly reduced the NAO-stained fluorescence in either wild-type or *metr-1(lf)*, indicating that methionine supplementation suppresses the increase of mitochondrial mass (Fig. 6B).

**Fig. 6.**
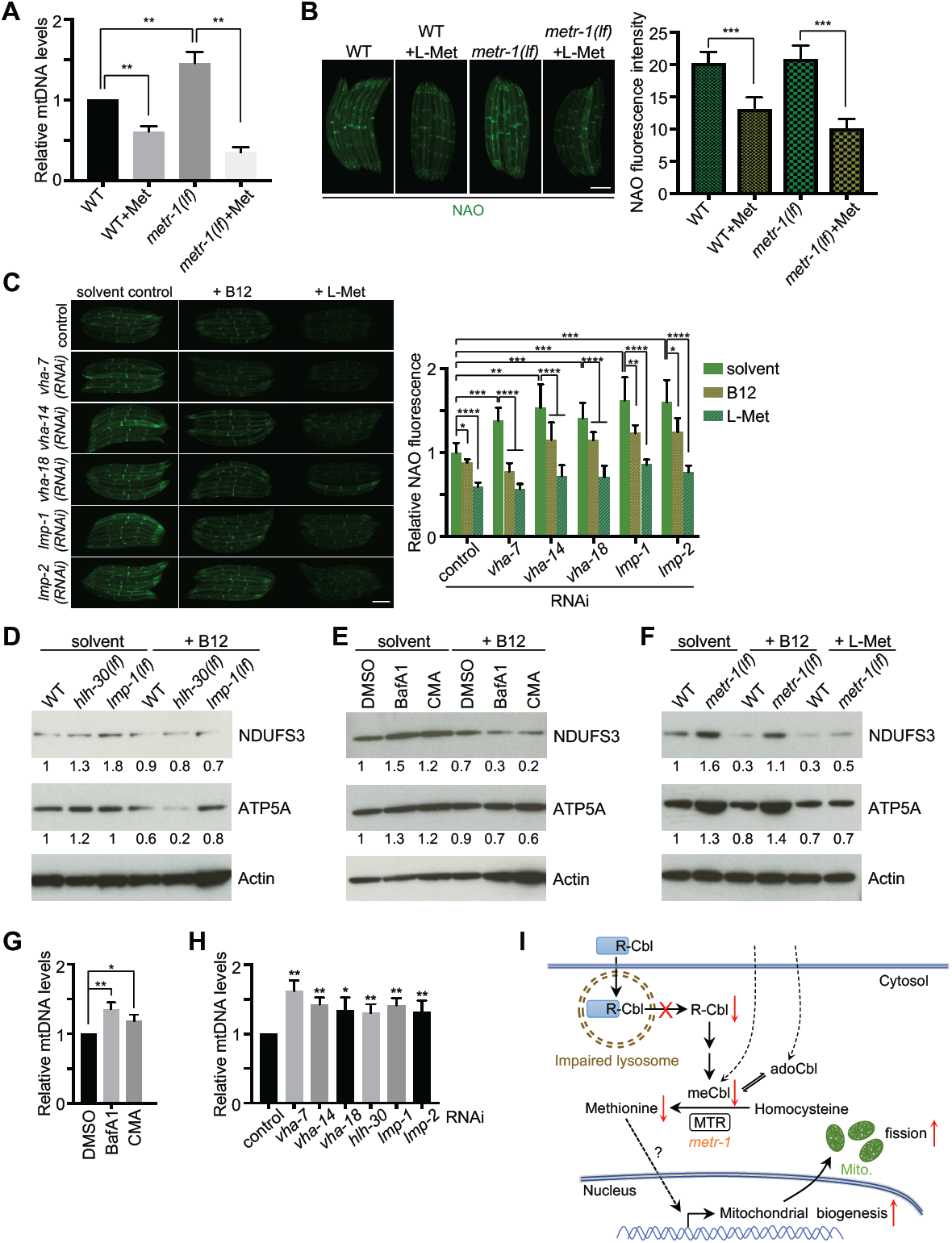
Lysosomal dysfunction leads to increase mitochondrial mass. (*A*) Relative mitochondrial DNA levels in wild-type and *metr-1(lf)* animals with or without methionine supplementation. Mean ± s.d. from at least three biological replicates. \*\**P* < 0.01. (*B* and *C*) Animals with indicated treatment were stained with Nonyl Acridine Orange (NAO). Mean ± s.d. \*\*\*\**P* < 0.0001, \*\*\**P* < 0.001, \*\**P* < 0.01, \*\**P* < 0.05. Scale bar, 0.2 mm. (*D*-*F*) Immunoblots of lysates from animals with indicated treatment. Relative protein levels were indicated below the gel lanes. (*G* and *H*) Relative mitochondrial DNA levels in animals with indicated treatment. Mean ± s.d. from at least three biological replicates. \*\**P* < 0.01, \**P* < 0.05. (*I*) A model for lysosomal activity affects mitochondrial fission through vitamin B12 metabolism.

We reasoned that lysosomal dysfunction caused B12 deficiency to then cause methionine restriction could lead to an increase of mitochondrial mass. Indeed, depletions of vacuolar V-ATPase subunits or lysosomal membrane proteins LMP-1 and LMP-2 caused a ∼ 1.5 fold increase in mitochondrial mass as measured by NAO staining (Fig. 6C). Moreover, supplementation of B12 or methionine suppressed the increase of NAO staining by lysosomal gene inactivations (Fig. 6C), suggesting that lysosomal dysfunction induced mitochondrial mass is mediated by B12 deficiency and methionine restriction. Consistent with these, detection of protein levels of conserved mitochondrial electron transport proteins using antibodies raised to highly conserved mammalian complex I component NDUFS3 and complex V component ATP5A showed increased mitochondrial mass in lysosomal mutants *hlh-30(lf)* and *lmp-1(lf)* (Fig. 6D), and in animals with V-ATPase inhibitor treatment (Fig. 6E), as well as in *metr-1(lf)* (Fig. 6F). B12 supplementation reduced these OXPHOS protein levels in all lysosomal defective animals as well as in wild-type, but not in *metr-1(lf)* (Figs. 6D-F). Supplementation of methionine readily reduced the protein levels of NDUFS3 and ATP5A in both wild-type and *metr-1(lf)* (Fig. 6F). These results indicate that lysosomal dysfunction increases mitochondrial mass mediated by B12 deficiency and methionine restriction. Moreover, inhibition of lysosomal function by V-ATPase inhibitor drugs or gene inactivations induced an increase in mtDNA copy number by ∼ 50% (Figs. 6G and 6H). In addition, RNAi of *hlh-30, lmp-1, lmp-2*, and V-ATPase subunit genes increased the expression of several mitochondrial biogenesis genes (Fig. S13A). These results suggest that lysosomal dysfunction increased mitochondrial biogenesis.

Mitophagy is selective macroautophagy that degrades damaged mitochondria. Lysosomal dysfunction could impair mitophagy and lead to accumulating damaged mitochondria to increase the mitochondrial abundance. However, we found that inactivations of the mitophagy pathway genes *pink-1, pdr-1* or *bec-1* did not increase, but in the cases of *pdr-1* and *bec-1* RNAi slightly decreased NAO staining in animals (Fig. S13B). In addition, inactivations of mitophagy genes did not change mtDNA levels (Fig. S13C), consistent with a previous observation (38). Moreover, mitophagy deficiency did not increase the expression of the tested mitochondrial biogenesis genes (Fig. S13D). Taking together, these results indicate that lysosomal dysfunction triggers an increase in mitochondrial mass by increasing mitochondrial biogenesis rather than by decreasing mitophagy.

## DISCUSSION

Mitochondrial dynamics is regulated by the balance of fission and fusion; imbalances in fission and fusion cause mitochondrial abnormalities that lead to many diseases, including neurodegeneration. From a screen of gene inactivations that suppress the mitochondrial fission defects of a *drp-1*/dynamin mutant, we identified a vacuolar V-ATPase subunit gene of the lysosome, *spe-5*. Testing other gene inactivations that cause lysosomal dysfunction showed that increased mitochondrial fission is a consequence of compromised lysosomal function. In yeast, age-dependent acidity decline in the lysosome-like vacuole is accompanied by mitochondrial fragmentation (39). In humans, lysosomal storage disorders (LSDs) are associated with mitochondrial fragmentation (40). These observations suggest a coupling between lysosomal function and mitochondrial fission across phylogeny. Our work further reveals that lysosomal dysfunction causes B12 deficiency by interrupting the release of vitamin B12 from the peroxisome. This B12 deficiency compromises the enzymatic activity of methionine synthase (MTR), a cytoplasmic enzyme that uses methylcobalamin as cofactor to convert homocysteine to methionine. Decreased methionine level, or methionine restriction, transduces the signal to the nucleus by yet to be identified factor(s) to augment mitochondrial biogenesis, which may be coupled to an increase of fission activity for new mitochondrial generation (Fig. 7I).

DRP-1/Drp1 dynamin mediates key cell biological steps in mitochondrial fission. However, our study shows that lysosomal dysfunction causes B12 deficiency and methionine restriction to increase mitochondrial fission, so that the fission-fusion cycles are rebalanced and mitigate the mitochondrial fission defect in a *drp-1*/dynamin mutant (Fig. S13E). These data show that there are pathways to mitochondrial fission that are independent of DRP-1 in *C. elegans*. Interestingly, previous studies also indicated that Drp1-independent fission exists in mammalian cells as mitochondria still undergo fragmentation in *Drp1*^*-/-*^ MEFs (41); and expression of a dominant negative mutant of Drp1, Drp1(K38A), could not completely inhibit hFis1-induced mitochondrial fragmentation (42). DRP-1/Drp1-independent fission, perhaps also via methionine biosynthetic pathways, may also operate in mammals.

The lysosome is required for carrier protein-mediated B12 transport and uptake in humans (34, 43) (Fig. 4F). Our data show that lysosomal dysfunction also causes vitamin B12 deficiency in *C. elegans* (Fig. S9). However, exogenous supplementation of metabolic active B12 reduced *hsp-6* induction and mitochondrial fragmentation defects even in a lysosomal mutation background (Figs. S10A and S10B), suggesting that B12 might be also absorbed by a carrier protein-independent/lysosome-independent method. As a small molecule, B12 may be passively transported, which is inefficient, but in our experiments we used a high dosage (13 nM is relatively high for a micronutrient such as vitamin B12). There is also passive transport for B12 in humans (44, 45).

B12 deficiency may disrupt the balance of mitochondrial dynamics under normal conditions (Fig. S13E). Mild B12 deficiency affected mitochondrial homeostasis and increased susceptibility to pathogenic infections in animals (46). However, in a *drp-1*/dynamin mutation background, B12 deficiency, on the contrary, improves *C. elegans* mitochondrial health by rebalancing mitochondrial fission-fusion events (Fig. S13E), demonstrating the elaborate regulation of mitochondrial dynamics.

B12 deficiency is common in the human population, particularly the elderly. Neurological disorders are significant clinical manifestations of B12 deficiency in humans (47-49). In yeast, lysosomal-like vacuolar acidity declines and mitochondria become gradually fragmented during aging (39). We observed that mitochondria become fragmented as *C. elegans* ages (data not shown). Increased mitochondrial fragmentation is widespread in lysosomal storage disorders (LSDs) as well as age-related neurodegeneration such as Alzheimer’s disease (AD), Parkinson’s disease (PD) and Huntington’s disease (HD) (50-53). Mutations affecting lysosomal function cause a class of metabolic diseases, or lysosomal storage disorders (LSDs), which are associated with neurologic symptoms and neurodegeneration (40). Mitochondrial abnormalities are a common feature for most LSDs (40), suggesting crosstalk between the two organelles. Based on our findings, we speculate that B12 deficiency caused by declines of lysosomal function may cause mitochondrial fragmentation during the aging process to in turn cause age-related neurological disorders and neurodegeneration. Dietary supplementation of B12 mitigated this mitochondrial fragmentation defect and the locomotion movement abnormalities in lysosomal defective animals (Fig. S10), suggesting the potential for vitamin B12 as a supplementary treatment in age-related neurological disorders and LSDs associated with mitochondrial fragmentation. Interference with the mitochondrial dynamics to re-balance the fission-fusion events could be a new therapeutic strategy for diseases with mitochondrial dysregulation.

## ACKNOWLEDGMENTS

We thank Bruce Bowerman for providing the *ruIs32*[P*pie-1*::GFP::H2B]III; *drp-1(or1393)IV* strain. Other strains were provided by the Caenorhabditis Genetics Center funded by NIH Office of Research Infrastructure Programs (P40 OD010440). We thank Ruvkun lab members for discussions and comments. This work was supported by an NIH grant to G.R. (NIH AG16636).

W.W. is supported by a Human Frontier Science Program Postdoctoral Fellowship.

## AUTHOR CONTRIBUTIONS

W.W. and G.R. designed experiments, analyzed data, and wrote the manuscript. W.W. performed all the experiments.

## FIGURE LEGENDS

**Fig. S1. RNAi screen identifies gene inactivation suppresses *drp-1* lethality**

(*A*) Schematic of the DRP-1 protein and the *drp-1(or1393*) and *drp-1(tm1108*) mutations.

(*B*) Diagram of the RNAi screen workflow for suppressors of *drp-1(or1393*) lethality. See Methods for details.

(*C*) Mitochondrial morphology in body wall muscles after indicated RNAi treatment. Scale bar, 5 μm.

(*D*) Viability of wild-type and *drp-1(or1393*) after indicated RNAi treatment at 23°C. Mean ±s.d. \*\*\**P* < 0.001, \*\**P* < 0.01.

(*E*) Viability of wild-type and *drp-1(tm1108*) in control versus *spe-5(RNAi*) animals. Mean ± s.d. \*\**P* < 0.01.

**Fig. S2. Representative images of mitochondrial morphology in indicated animals**

Mitochondrial morphology of wild-type and *drp-1(tm1108*) in control versus *spe-5(RNAi*) in a single body wall muscle cell. Mitochondria are visualized by a mitochondrial outer membrane localized mRFP fusion protein. Lower panels are plot profiles of the cross sections indicated by yellow lines. Criterion for a peak: peak to trough (both sides) > 20 Grey value units (y units). Scale bar, 5 μm.

**Fig. S3. Inactivation of the vacuolar ATPase *spe-5* suppresses *drp-1* mitochondrial defects**

(*A*) P*hsp-6*::GFP expression in wild-type and *drp-1(tm1108*) animals grown on control versus *spe-5(RNAi*). Mean ± s.d. \*\*\**P* < 0.001, \**P* < 0.05. Scale bar, 0.2 mm.

(*B*) Endogenous expression levels of mitochondrial chaperone genes *hsp-6* and *hsp-60* in indicated animals. Mean ± s.d. from at least three biological replicates. \*\*\**P* < 0.001, \*\**P* < 0.01, \**P* < 0.05.

(*C*) Animals with indicated treatment were stained with TMRE (tetramethylrhodamine, ethyl ester). Mean ± s.d. \*\*\**P* < 0.001. Scale bar, 0.2 mm.

**Fig. S4. Disruptions of V-ATPase activity suppress *drp-1* lethality and mitochondrial defects**

(*A, B*) Viability of wild-type and *drp-1(or1393*) with DMSO, BafA1 or CMA treatment at 23°C. DMSO, solvent control; BafA1, bafilomycin A1; CMA, concanamycin A. Mean ± s.d. \*\**P* < 0.01. Scale bar, 1 mm.

(*C, D*) P*hsp-6*::GFP expression in wild-type and *drp-1(tm1108*) animals with indicated treatment. Mean ± s.d. \*\*\*\**P* < 0.0001, \*\**P* < 0.01, \**P* < 0.05. n.s., not significant. Scale bar, 0.2 mm.

(*E, F*) Endogenous expression levels of mitochondrial chaperone genes *hsp-6* and *hsp-60* in animals with indicated treatment. Mean ± s.d. from at least three biological replicates. \*\*\*\**P* < 0.0001, \*\*\**P* < 0.001, \*\**P* < 0.01, \**P* < 0.05.

**Fig. S5. Inhibitions of V-ATPase activity increase mitochondrial membrane potential in *drp-1(tm1108*)**

(*A, B*) Animals with indicated treatment were stained with TMRE. Mean ± s.d. \*\*\*\**P* < 0.0001, \*\**P* < 0.01. Scale bar, 0.4 mm.

**Fig. S6. Lysosomal dysfunction suppresses *drp-1* mitochondrial fission defects**

(*A*) P*hsp-6*::GFP expression in wild-type and *drp-1(tm1108*) animals grown on control or indicated RNAi bacteria clones. Mean ± s.d. \*\**P* < 0.01. n.s., not significant. Scale bar, 0.2 mm.

(*B*) Endogenous expression levels of mitochondrial chaperone genes *hsp-6* and *hsp-60* in animals with indicated treatment. Mean ± s.d. from at least three biological replicates. \*\*\*\**P* < 0.0001, \*\*\**P* < 0.001, \*\**P* < 0.01.

(*C*) Animals with indicated treatment were stained with TMRE. Mean ± s.d. \*\*\**P* < 0.001, \*\**P* < 0.01. Scale bar, 0.2 mm.

(*D*) Mitochondrial morphology in *drp-1(tm1108*) body wall muscles after indicated RNAi treatment. Scale bar, 5 μm.

(*E*) Percentage of mitochondria with tubular morphology in *drp-1(tm1108*) body wall muscles after indicated RNAi treatments. Mean ± s.d.

(*F*) Viability of *drp-1(or1393*) after mitophagy gene inactivations at 23°C.

**Fig. S7. Dietary B12 deficiency mitigates *drp-1* defects**

(*A*) Growth of wild-type (WT) and *drp-1(tm1108*) fed on *E. coli* HT115 or *E. coli* OP50 with or without B12 supplementation at 20°C for 5-day since embryos. Scale bar, 1 mm.

(*B*) P*hsp-6*::GFP expression and animal lengths in wild-type and *drp-1(lf*) animals raised on *E. coli* HT115 or *E. coli* OP50 with or without B12 supplementation. Mean ± s.d. \*\*\**P* < 0.001, \*\**P* < 0.01. *drp-1(lf), drp-1(tm1108*). Scale bar, 0.2 mm.

**Fig. S8. P*acdh-1*::GFP as vitamin B12 indicator**

(*A*) P*acdh-1*::GFP animals raised on live (upper panels) or dead (75°C 0.5h, lower panels) *E. coli* HT115 or *E. coli* OP50 with or without B12 supplementation. Scale bar, 0.2 mm.

It is possible that *E. coli* K12 and *E. coli* B strains metabolize B12 differently to affect the animals grown on them. To exclude this possibility, we killed the *E. coli* bacteria by heat. Incubation of the *E. coli* bacteria at 70°C for 30 min was enough to completely kill them.

Animals fed on dead bacteria induced P*acdh-1*::GFP on *E. coli* OP50 (B12 deficient) diets, although much milder than those fed on live bacteria.

(*B*) Bacteria with or without heat treatment grown on LB agar plate at 37°C overnight.

(*C*) P*acdh-1*::GFP expression for animals raised on live *E. coli* HT115 or *E. coli* OP50 with or without B12 supplementation. Mean ± s.d. \*\*\*\**P* < 0.0001.

(*D*) P*acdh-1*::GFP expression for animals raised on *E. coli* OP50 without or with different doses of vitamin B12 supplementation. Scale bar, 0.2 mm.

(*E*) P*acdh-1*::GFP expression for animals raised on *E. coli* OP50 with or without propionic acid treatment and B12 supplementation. Propionic acid treatment induced P*acdh-1*::GFP, which was suppressed by vitamin B12 supplementation. Scale bar, 0.2 mm.

**Fig. S9. Lysosomal dysfunction leads to B12 deficiency**

(*A, B*) P*acdh-1*::GFP expression for animals with indicated treatment. Mean ± s.d. \*\*\*\**P* < 0.0001, \*\*\**P* < 0.001, \*\**P* < 0.01, \**P* < 0.05. Scale bar, 0.2 mm.

(*C*) P*acdh-1*::GFP expression for animals with indicated genetic mutation background. Mean ±s.d. \*\*\*\**P* < 0.0001, \*\*\**P* < 0.001, \*\**P* < 0.01. Scale bar, 0.1 mm.

(*D*) Survival of animals treated with solvent control, 100 mM propionic acid, or 15 mM homocysteine (Hcy). Mean ± s.d. \*\*\**P* < 0.001, \*\**P* < 0.01.

Although the propionate shunt is activated when the cellular B12 level is low to breakdown the excess propionate, high propionate loading saturates the propionate shunt. Indeed, a *C. elegans mmcm-1* loss-of-function mutation reduced the survival of animals in the presence of 100 mM propionic acid, consistent with the finding that deletion of PCCA-1, an upstream enzyme of the canonical propionate breakdown pathway, reduced the LD_50_ for propionate (54). The *metr-1* methionine synthetase loss-of-function mutation caused animals to be resistant to a high concentration of propionate on an *E. coli* OP50 diet, similar to previous observation (27). Deletion of either *mmcm-1* or *metr-1* caused animals to become hypersensitive to exogenously supplementation of 15 mM homocysteine. Both *hlh-30(lf*) and *lmp-1(lf*) lysosomal mutants were hypersensitive to 100 mM propionic acid and 15 mM homocysteine compared to wild-type, suggesting B12 deficiency in these lysosomal mutants. Overall, these findings indicate that lysosomal dysfunction causes B12 deficiency in animals that were fed B12 proficient *E. coli* HT115 K12 strains, which usually provide enough vitamin B12 to wild-type animals.

**Fig. S10. Dietary B12 supplementation rescues mitochondrial fragmentation and locomotion defects caused by lysosomal dysfunction**

(*A) lmp-1(lf*); P*hsp-6*::GFP animals fed on *E. coli* HT115 or *E. coli* OP50 with or without B12 supplementation. Mean ± s.d. \*\*\**P* < 0.001, \*\**P* < 0.01. Scale bar, 0.1 mm.

(*B*) Mitochondrial morphologies and lengths in body wall muscles in *lmp-1(lf*) fed on *E. coli* HT115 or *E. coli* OP50 with or without B12 supplementation. Arrows mark some examples of fragmented mitochondria. Mitochondrial lengths were calculated by MiNA toolset. Median with 95% C. I. Mann-Whitney test. \*\*\*\**P* < 0.0001. N indicates the sample size. Scale bar, 5 μm. (*C, D*) Locomotion movement of wild-type (WT) or *lmp-1(lf*) fed on *E. coli* HT115 or *E. coli* OP50 with or without B12 supplementation. n=25∼40. \*\*\**P* < 0.001.

**Fig. S11. Methionine restriction increases mitochondrial fission**

(*A*) Development of wild-type, *lmp-1(lf*) or *drp-1(lf*) grown on indicated RNAi clones. Scale bar, 1 mm

(*B*) P*hsp-6*::GFP with wild-type, *lmp-1(lf*) and *drp-1(lf*) background animals grown on indicated RNAi clones. Mean ± s.d. \*\*\**P* < 0.001, \*\**P* < 0.01. n.s., not significant. Scale bar, 0.2 mm.

(*C*) P*hsp-6*::GFP expression in *lmp-1(lf*) (upper panels) and wild-type (lower panels) animals grown on indicated bacteria with or without methionine supplementation. Mean ± s.d. \*\*\**P* < 0.001, \*\**P* < 0.01. Scale bar, 0.1 mm.

**Fig. S12. Inactivation of *sams-1* increases mitochondrial fission**

(*A*) Mitochondrial morphology in wild-type and *drp-1(lf*) body wall muscles after indicated RNAi treatment. Scale bar, 5 μm.

(*B*) P*hsp-6*::GFP animals grown on control versus *sams-1(RNAi*). Scale bar, 0.2 mm.

(*C*) Mitochondrial morphology in animals grown on control or *sams-1(RNAi*) after indicated treatment. Scale bar, 5 μm.

**Fig. S13. Lysosomal dysfunction increases mitochondrial mass by inducing mitochondrial biogenesis**

(*A, D*) Expression levels of mitochondrial biogenesis genes *atp-5, cox-4 hmg-5* and *gas-1* in animals after indicated RNAi treatment. Mean ± s.d. from at least three biological replicates.

(*B*) Animals with mitophagy gene inactivations were stained with Nonyl Acridine Orange (NAO). Mean ± s.d. \*\*\*\**P* < 0.0001, \*\**P* < 0.01. Scale bar, 0.2 mm.

(*C*) Relative mitochondrial DNA levels in control and mitophagy defective animals. Mean ± s.d. from at least three biological replicates. n.s., not significant.

(*E*) The schematic diagram of the balance of mitochondrial fission-fusion events under different physiological condition.

## METHODS

### *C. elegans* strains and maintenance

*C. elegans* were grown on standard nematode growth medium at 20 °C, or other temperatures when indicated. To synchronize animals, embryos were harvested from gravid hermaphrodites treated with bleach solution containing 0.5 M NaOH, allowed to hatch in M9 buffer overnight. The following strains were used in this study: N2 Bristol: wild-type, CU6372: *drp-1(tm1108)IV*, RB2375: *lmp-1(ok3228)X*, JIN1375: *hlh-30(tm1978)IV*, RB755: *metr-1(ok521)II*, RB1434: *mmcm-1(ok1637)III*, PS6192: *syIs243*[P*myo-3*::TOM20::mRFP], SJ4100: *zcIs13*[P*hsp-6*::GFP]V, VL749: *wwIs24*[P*acdh-1*::GFP]. *drp-1(or1393)IV* was generated by out-crossing EU2706: *ruIs32*[P*pie-1*::GFP::H2B]III; *drp-1(or1393)IV* provided by B. Bowerman into N2. *drp-1(lf*); P*myo-3*::TOM20::mRFP was generated by crossing *drp-1(tm1108*) into P*myo-3*::TOM20::mRFP. *drp-1(lf*); P*hsp-6*::GFP was generated by crossing *drp-1(tm1108*) into P*hsp-6*::GFP. *lmp-1(lf*); P*myo-3*::TOM20::mRFP was generated by crossing *lmp-1(ok3228*) into P*myo-3*::TOM20::mRFP. *lmp-1(lf*); P*hsp-6*::GFP was generated by crossing *lmp-1(ok3228*) into P*hsp-6*::GFP.

### RNA interference

*E. coli* HT115 bacteria expressing targeted double-strand RNAs were cultured in LB containing 50 μg/ml ampicillin at 37 °C overnight and seeded 150 μl per 60 mm or 100 μl per well of 24-well RNAi plate containing 5 mM IPTG and 100 μg/ml carbenicillin. Plates were allowed to dry and induce at room temperature overnight. Synchronized animals were transferred to the RNAi plates and cultured at 20 °C or at indicated temperatures depending on the experiments. For all RNAi experiments, an empty L4440 vector was used as the control.

For RNAi screen for *drp-1* suppressors, ∼15 synchronized L1 *drp-1(or1393*) were dropped on each well on 24-well RNAi plates and kept at 23 °C for 3 d when the animals laid plenty of eggs. Continued to the culture at 23 °C for another 48 h and then scored the hatched progeny on each well.

### Viability assays

Synchronized L1 animals were grown on indicated RNAi plates at 20 °C, 23 °C or 25 °C for 3 d, or at 15 °C for 5 d when they reached adults. ∼ 20 gravid adult animals were transferred to new RNAi plates and allowed to lay eggs for 2 h and then removed. Egg numbers were counted. The eggs were continued to keep at corresponding temperatures for another 48 h. Hatched progeny was then scored for viability. Bright field Images represented the viability were obtained by a Zeiss AxioZoom V16 microscope.

### Mitochondrial morphology and length

After indicated treatment, synchronized adult animals containing the P*myo-3*::TOM20::mRFP transgene were mounted on 2% agarose pads for microscopic imaging. Each treatment scored mitochondrial morphology in 8∼10 body wall muscles in the middle of the worm body and 10∼15 animals in total. Mitochondria exhibited tubular structure in most parts of a single muscle cell were defined as ones with “tubular morphology”, while exhibited elongated structure in most parts of a muscle was defined as ones with “elongated morphology”, and “fragmented morphology” indicated that mitochondria in a muscle cell displayed many fragmented pieces. Fluorescence images represented mitochondrial morphology were obtained by a Zeiss Axio Image Z1 microscope.

Mitochondrial length was quantified by ImageJ software as described (Valente et al., 2017) with modification. Briefly, the images were preprocessed (Unsharp mask → Enhance local contrast → Median), then converted to binary and generated a morphological skeleton for calculating the “branch lengths” as the mitochondrial lengths. Statistical analyses with Mann-Whitney test were performed with Prism 7.

### Induction of GFP reporters

Synchronized L1 animals were grown on indicated culture plates at 20 °C for 3-4 d. Dropped ∼ 2 μl of 5 mM levamisole (Sigma) onto a new NGM plate and transferred several animals into it. Images were captured by a Zeiss AxioZoom V16 microscope with the same exposure time and magnification. Fluorescence intensities or pixel intensities were measured by ImageJ software. Statistical analyses were done from at least 3 biological replicates.

### Chemical treatment

To inhibit V-ATPase activity, synchronized L1 animals were grown at 20 °C for 2 d until they reached late L4s. 100 μl M9 buffer containing 10 μg/ml Bafilomycin A1 (LC Laboratories) dissolved in DMSO or 1 μg/ml concanamycin A (AG Scientific, Inc) dissolved in DMSO was directly added onto the bacterial lawn. Plates were air dried and kept at 20 °C for another 48 h.

For B12 supplementation, animals were directly grown on culture plates containing 12.8 nM adenosylcobalamin (adoCbl) or methylcobalamin (meCbl) (Sigma). For methionine supplementation, animals were grown on culture plates containing 13.4 mM L-methionine (Sigma).

For propionic acid and homocysteine hypersensitivity assays, ∼ 100 embryos for each genetic mutation were transferred to culture plates containing solvent control, 100 mM propionic acid (Sigma) or 15 mM homocysteine (Sigma). 2 days later scored the hatched animals that had developed into L3 or later stages. Statistical analyses were done from at least 3 biological replicates.

### NAO and TMRE staining

Synchronized L1 N2 were grown on indicated plates at 20 °C for 2 d until they reached late L4. 5 μM Nonyl Acridine Orange (Acridine Orange 10-Nonyl Bromide) (Life Technologies) or 1 μM TMRE (tetramethylrhodamine, ethyl ester) (Life Technologies) in 100 μl M9 buffer was then directly added onto the bacterial lawn. Plates were air dried and kept at 20 °C overnight. Animals were then picked and transferred to new RNAi plates without Nonyl Acridine Orange or TMRE to grow for 8 h to clean any non-uptake fluorescent dye. Images were then obtained with a Zeiss AxioZoom V16 microscope using the same exposure time and magnification and then processed by ImageJ software.

### mtDNA levels

Synchronized L1 N2 were grown on standard NGM plates seeded with *Escherichia coli* OP50 at 20 °C for 2 d until they reached late L4s, then incubated with desired drugs or transferred to the desired RNAi plates and kept at 20 °C for another 48 h. Animals were then harvested and total DNA was extracted and quantified using quantitative real-time PCR as described previously (55). Primers were designed to quantify *nd1* (for mitochondrial DNA) and *act-3* (for genomic DNA) as previously described (56), and mitochondrial DNA levels were then normalized to genomic DNA.

### RNA isolation and quantitative RT-PCR

Synchronized L1 animals were grown on standard NGM plates seeded with *Escherichia coli* OP50 at 20 °C for 2 d until they reached late L4s, then incubated with desired drugs or transferred to the desired RNAi plates and kept at 20 °C for another 48 h. Animals were then harvested and total RNA was extracted by RNeasy Plus Universal Kit (Qiagen) following the manufacturer’s instructions. 1 μg total RNA was then used for first strand cDNA synthesis using QuantiTect Reverse Transcription Kit (Qiagen). Quantitative real-time PCR was performed using iQ SYBR Green Supermix (Bio-Rad). Quantification of transcripts was normalized to *act-4* (57) and relative mRNA levels were calculated by ΔΔCt. Primers for desired targets were designed to span exon/exon boundaries.

*atp-5*: 5’-ACGTCTTCCGTGAACTGGTCGAAG-3’ and 5’-GCAAAGTCGATCTTGGGAAGATCG-3’; *cox-4*: 5’-TCGTCGGATACGGAGCAAATG-3’ and 5’-TCGAATTCGGACAGCGTTTGAC-3’; *hmg-5*: 5’-GAAGTTGTCTGGAGCTGGAATG-3’ and 5’-GGAGGACGACATGGTATTCATC-3’; *gas-1*: 5’-CAATTGAGGCTCCAAAGGGAGAG-3’ and 5’-GACATGTAGCACACATCGTGGATG-3’; *gst-4*: 5’-GTCTATCACAAGATACTTGGCAAG-3’ and 5’-ATCACGGGCTGGTTCAACAACTTC-3’; gst-10: 5’-TGCAGACTGGAGCAATTATGCGTC-3’ and 5’-TCCCTCGTCGTAGTAAATGTAACG-3’.

### Western blotting and Antibodies

Synchronized L1 animals were grown on plates with indicated treatment at 20 °C for 3 d. Animals were then harvested and washed with M9 buffer for 2∼3 times, then resuspended with 1X NuPAGE LDS Sample Buffer (Invitrogen) containing 5% β-mercaptoethanol and heated at 70 °C for 10 min. Lysates were loaded onto NuPAGE 4-12% Bis-Tris Protein Gels (Invitrogen) and transferred onto Nitrocellulose membrane (Invitrogen). The membrane was then blocked with 5% nonfat milk and probed with designated primary and secondary antibodies.

The primary antibodies used in this study included anti-NDUFS3 (Abcam, ab14711), anti-ATP5A (Abcam, ab14748), and anti-Actin (Abcam, ab179467).

### Statistical analysis

No statistical methods were used to predetermine sample sizes. Statistical tests with appropriate underlying assumptions on data distribution and variance characteristics were used. All samples were randomly selected and from at least 3 biological replicates when applied. Data were analyzed with one-factor ANOVA (for mitochondrial morphology experiments) and two-tailed unpaired Student’s *t*-tests (for the other experiments). All statistical analyses and graphing were performed with Prism 7 (GraphPad software). Statistically significant differences were indicated as * *P* < 0.05, ** *P* < 0.01, *** *P* < 0.001, **** *P* < 0.0001, and n.s. as not significant.

