## Supplemental Figures for "Lysosomal activity regulates *Caenorhabditis elegans* mitochondrial dynamics through vitamin B12 metabolism"

### Supplementary Figure 1

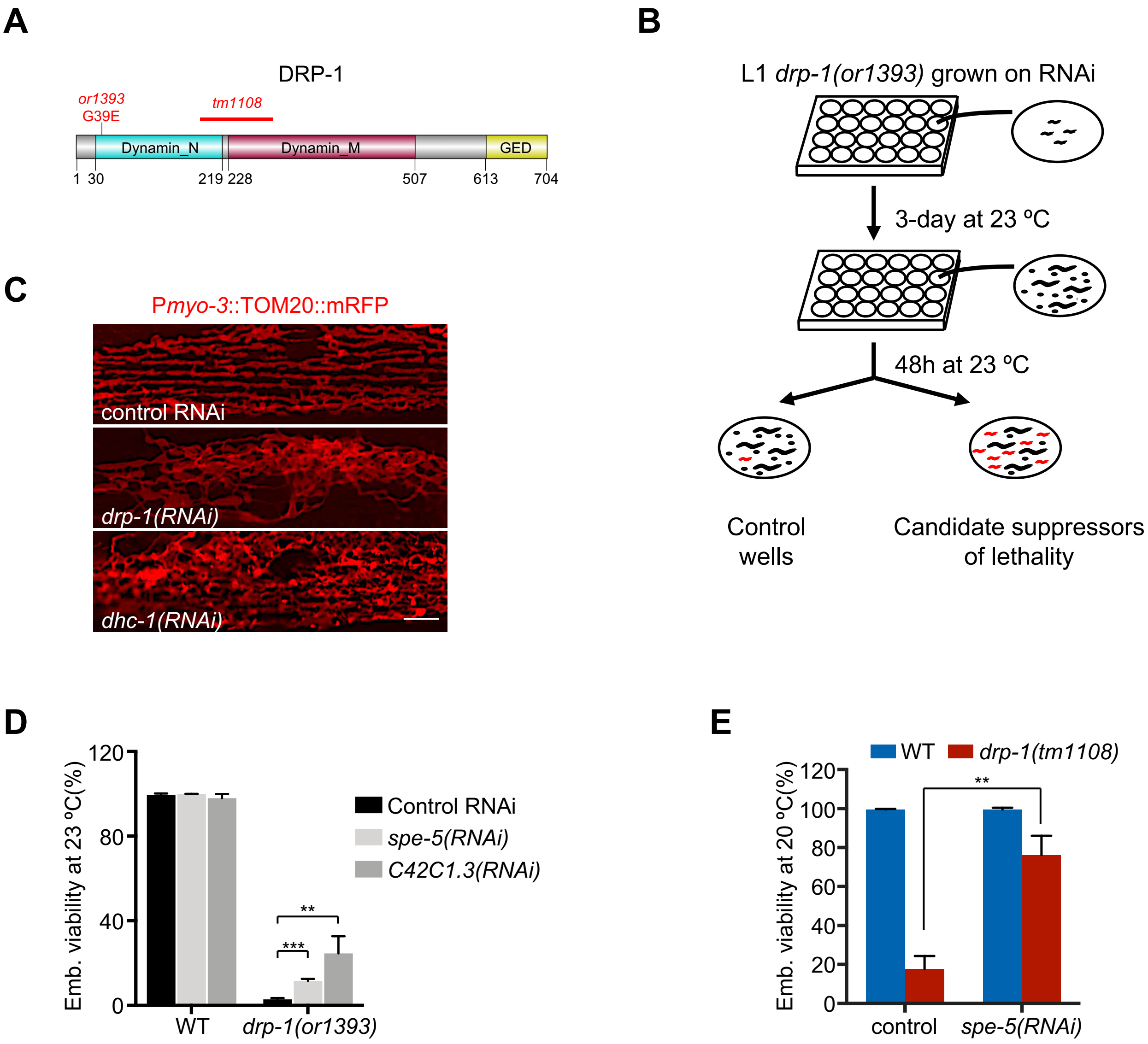

### Supplementary Figure 2

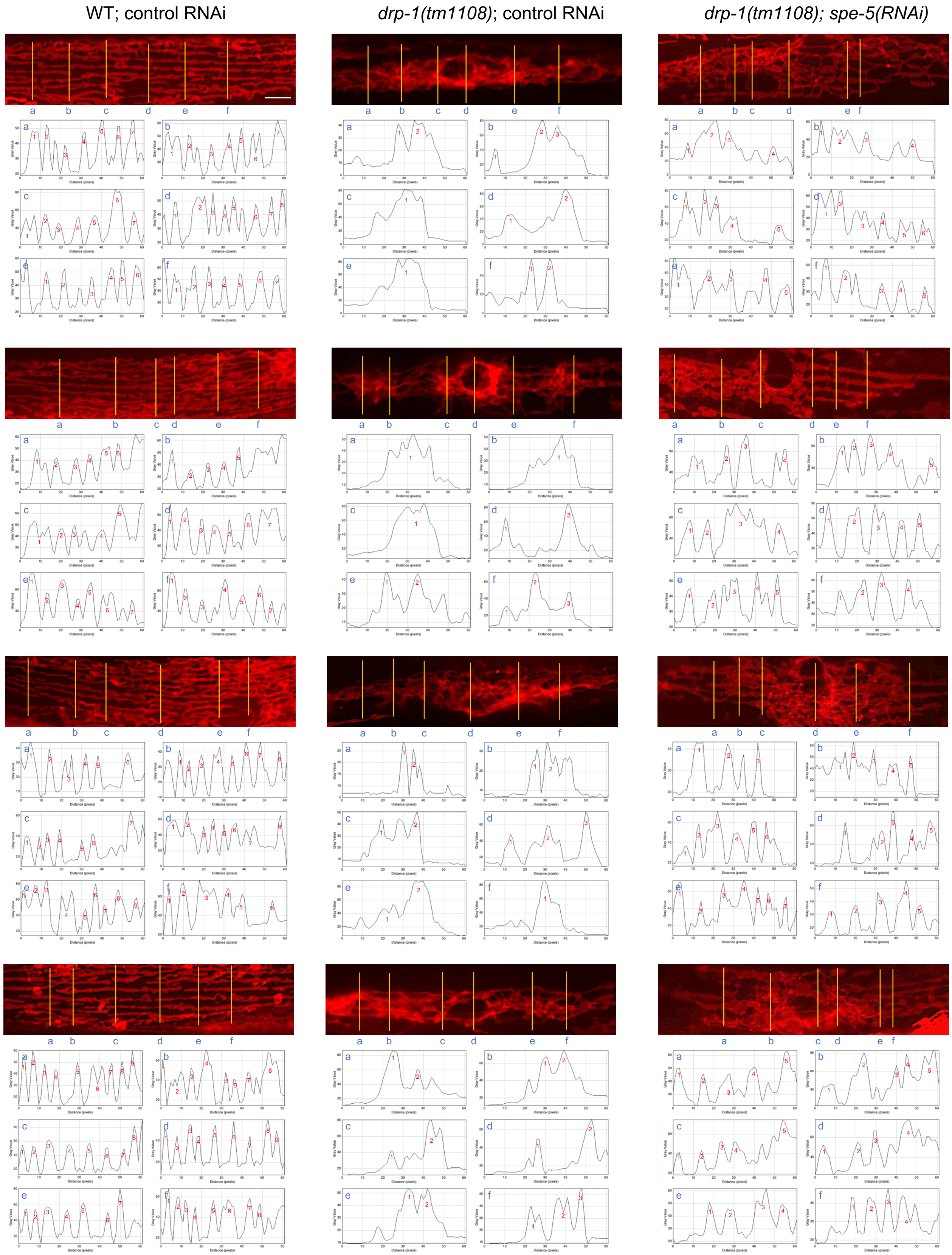

### Supplementary Figure 3

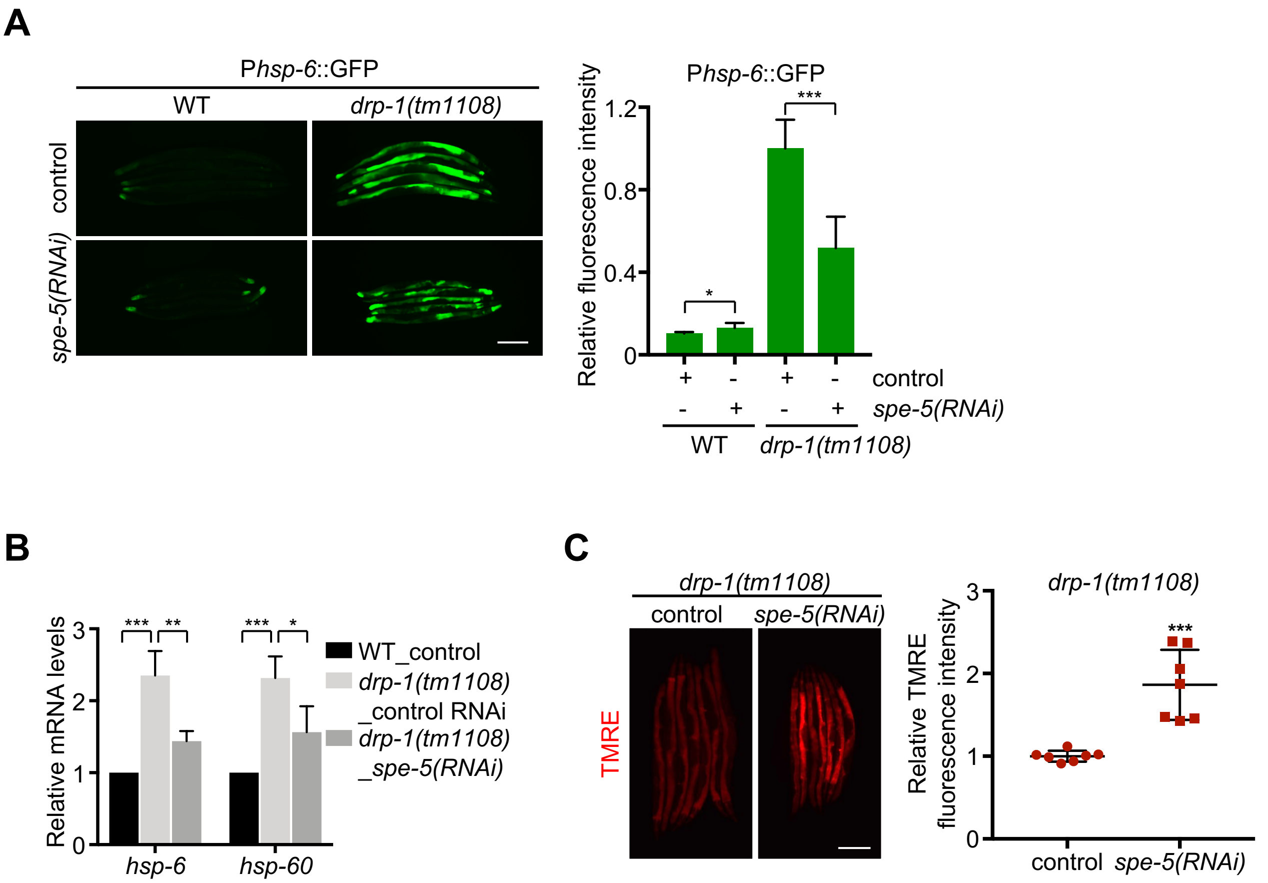

### Supplementary Figure 4

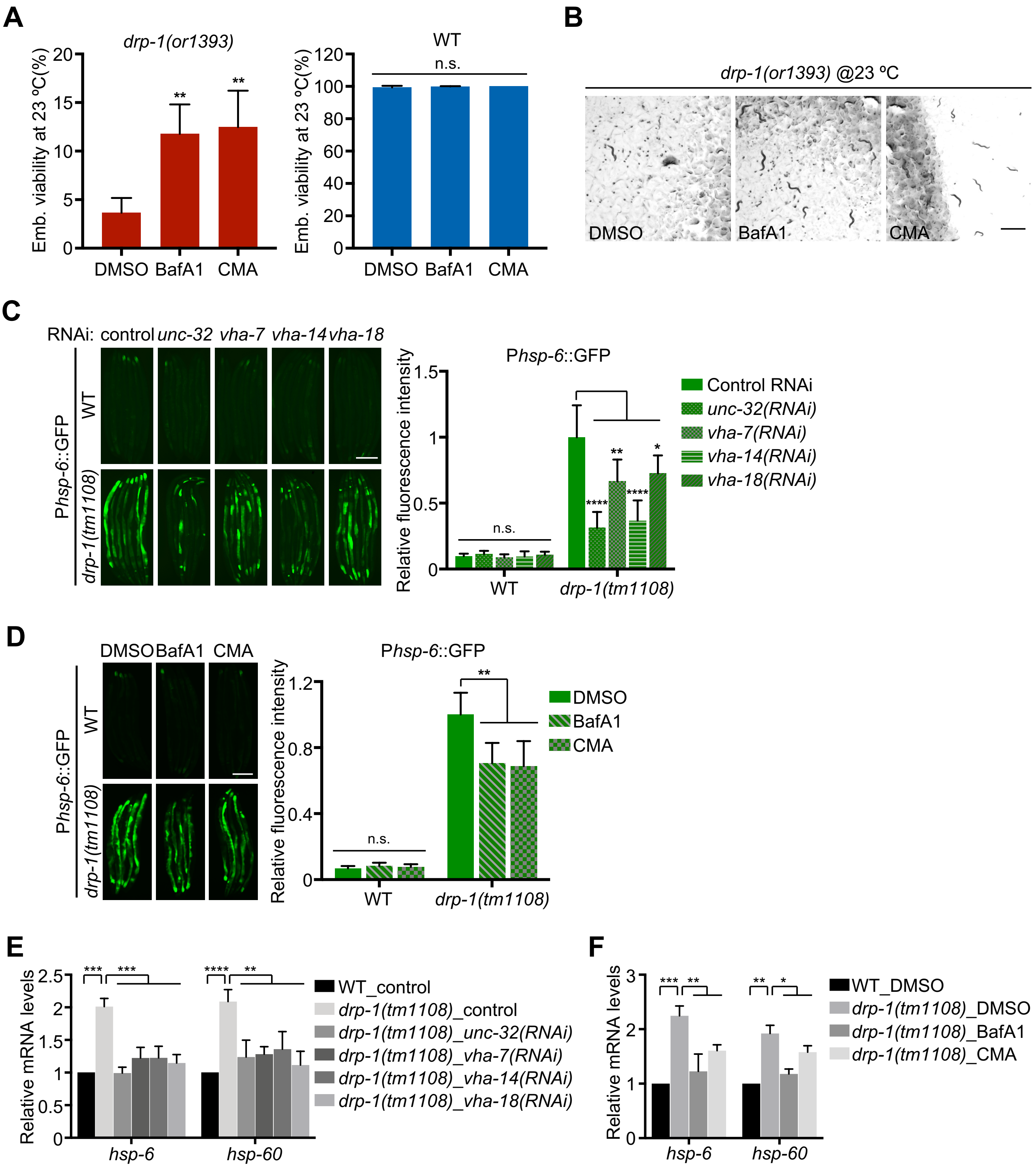

### Supplementary Figure 5

A

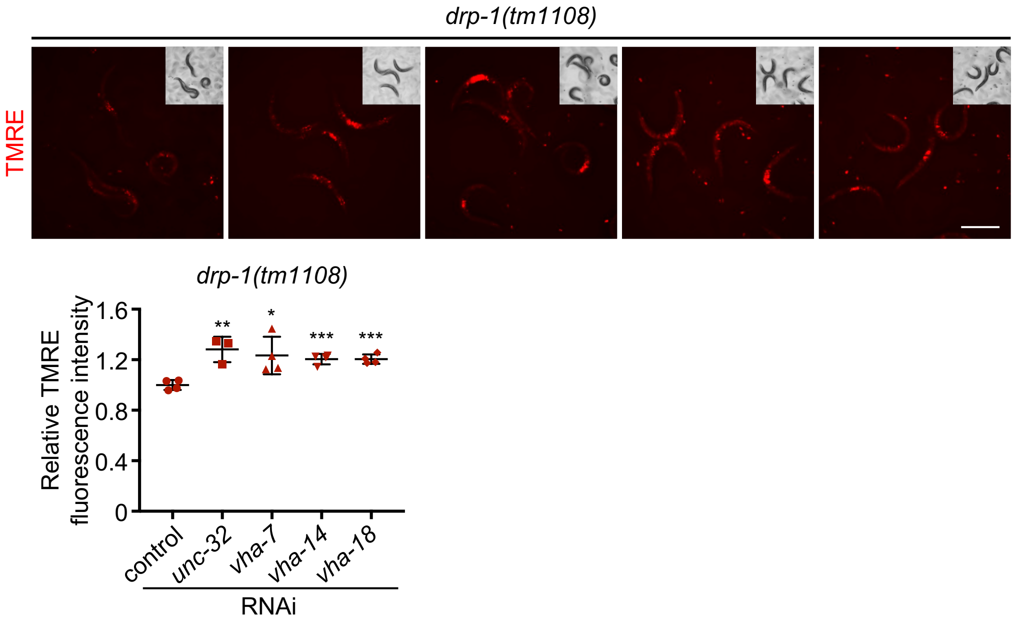


B

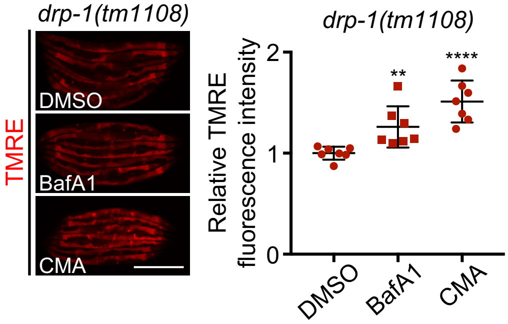

#### Supplementary Figure 6

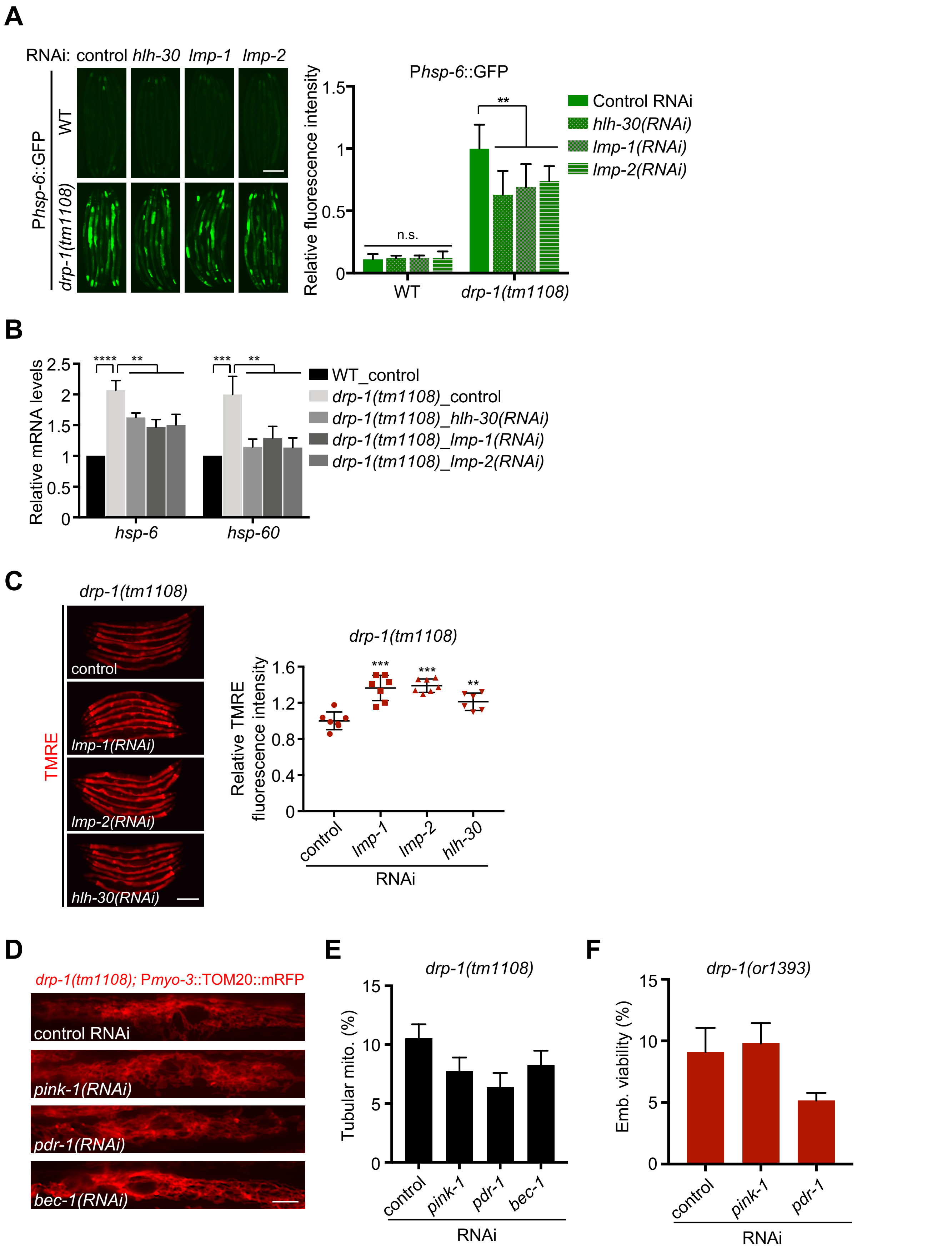

### Supplementary Figure 7

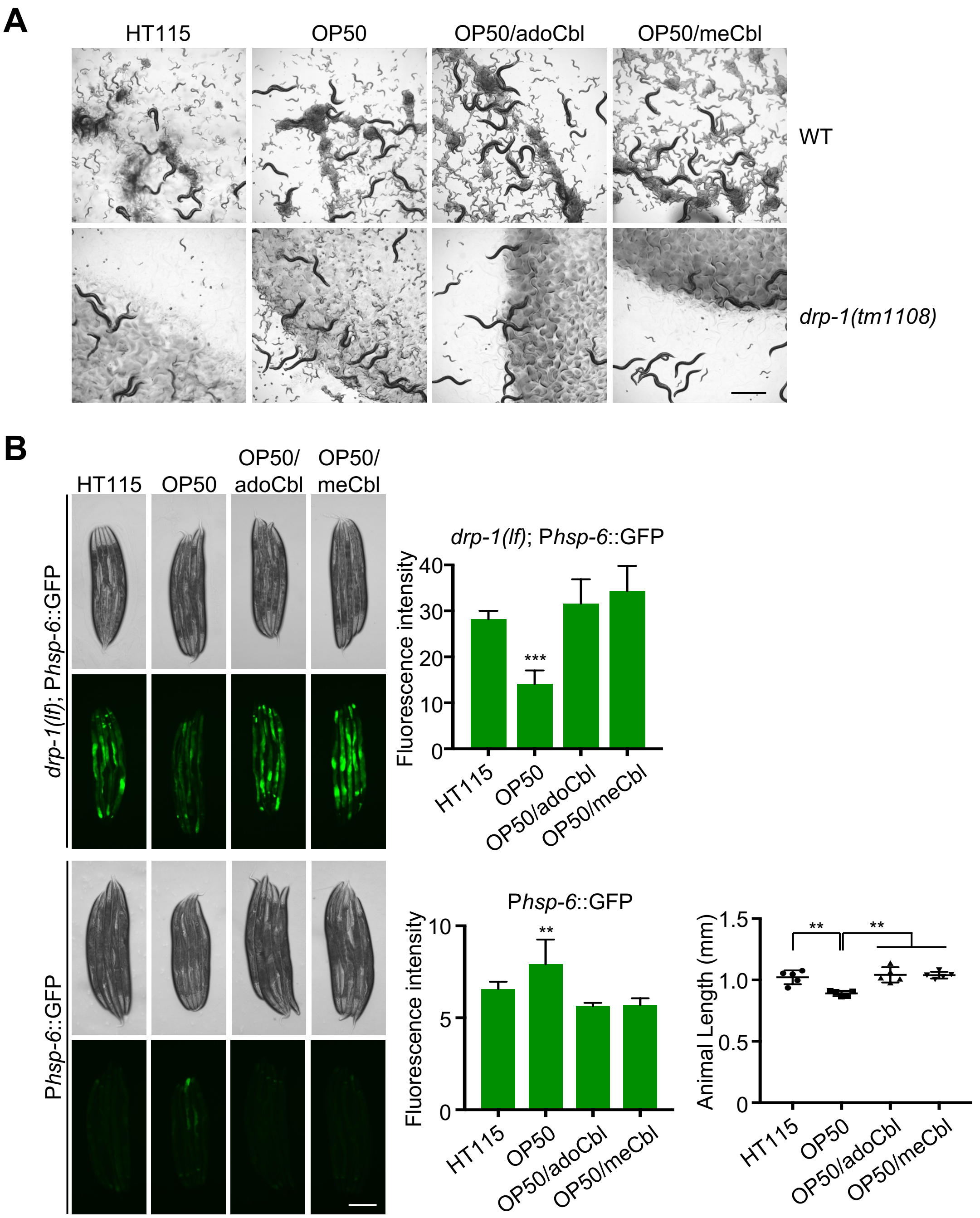

### Supplementary Figure 8

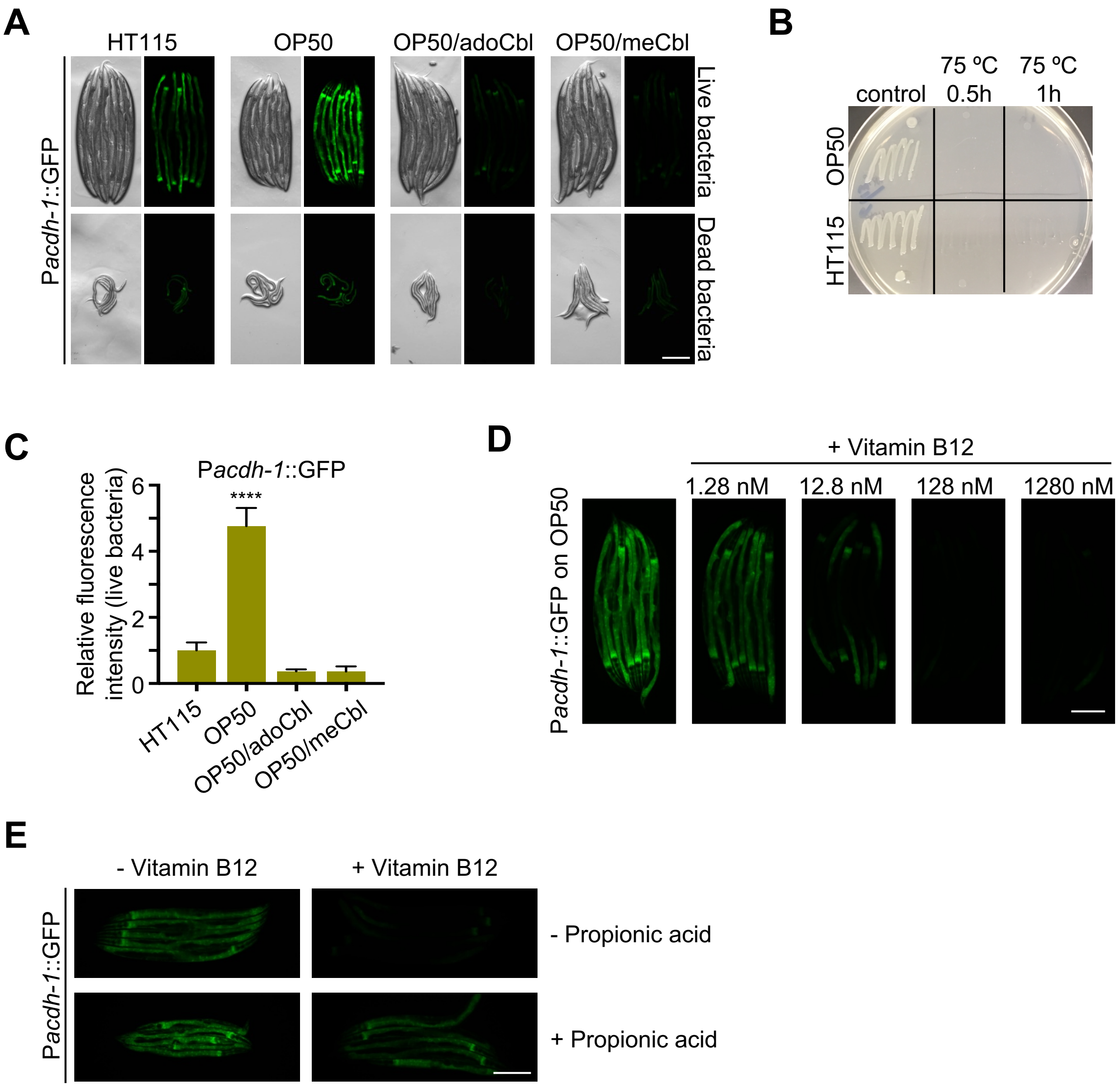

### Supplementary Figure 9

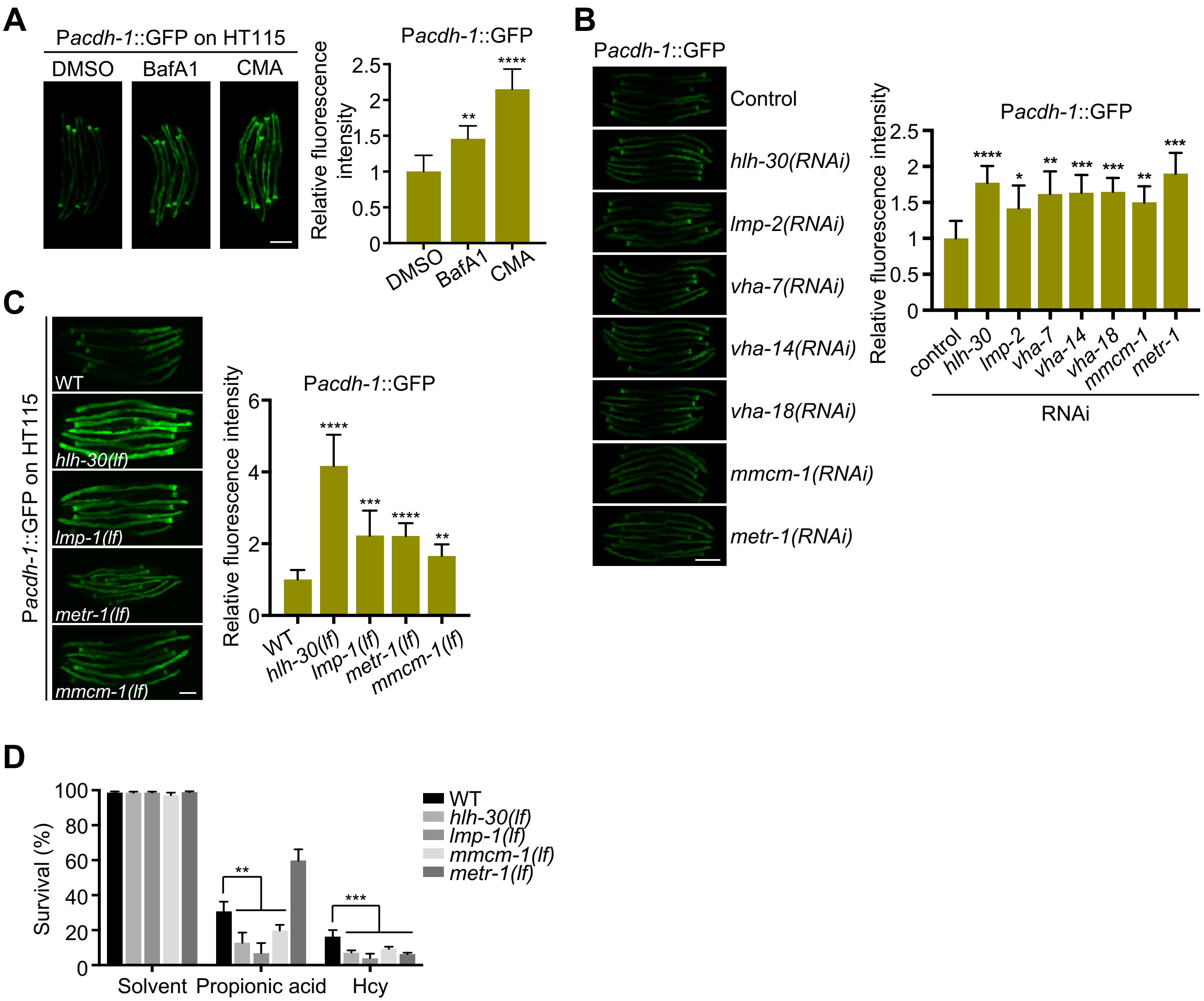

### Supplementary Figure 10

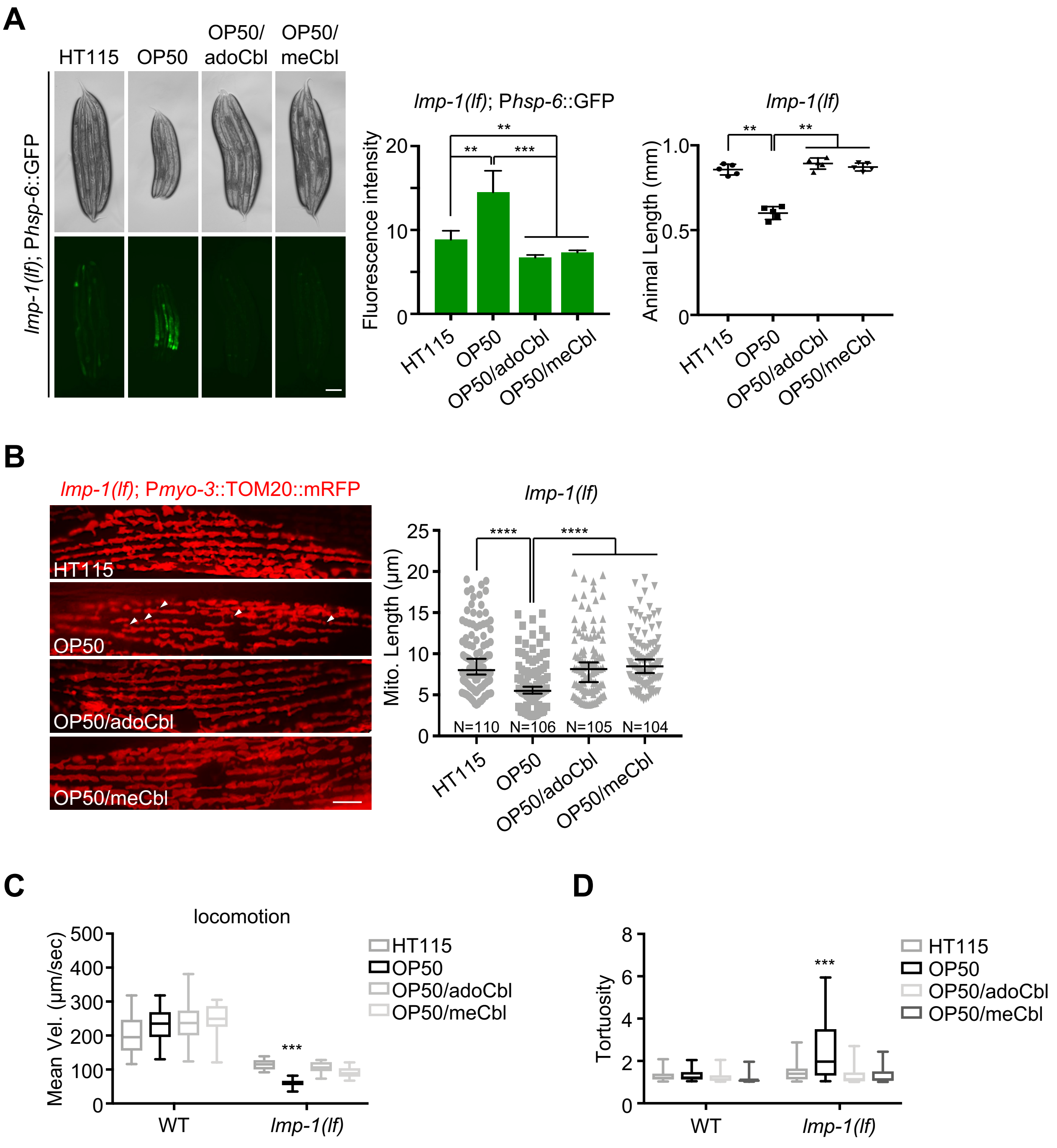

### Supplementary Figure 11

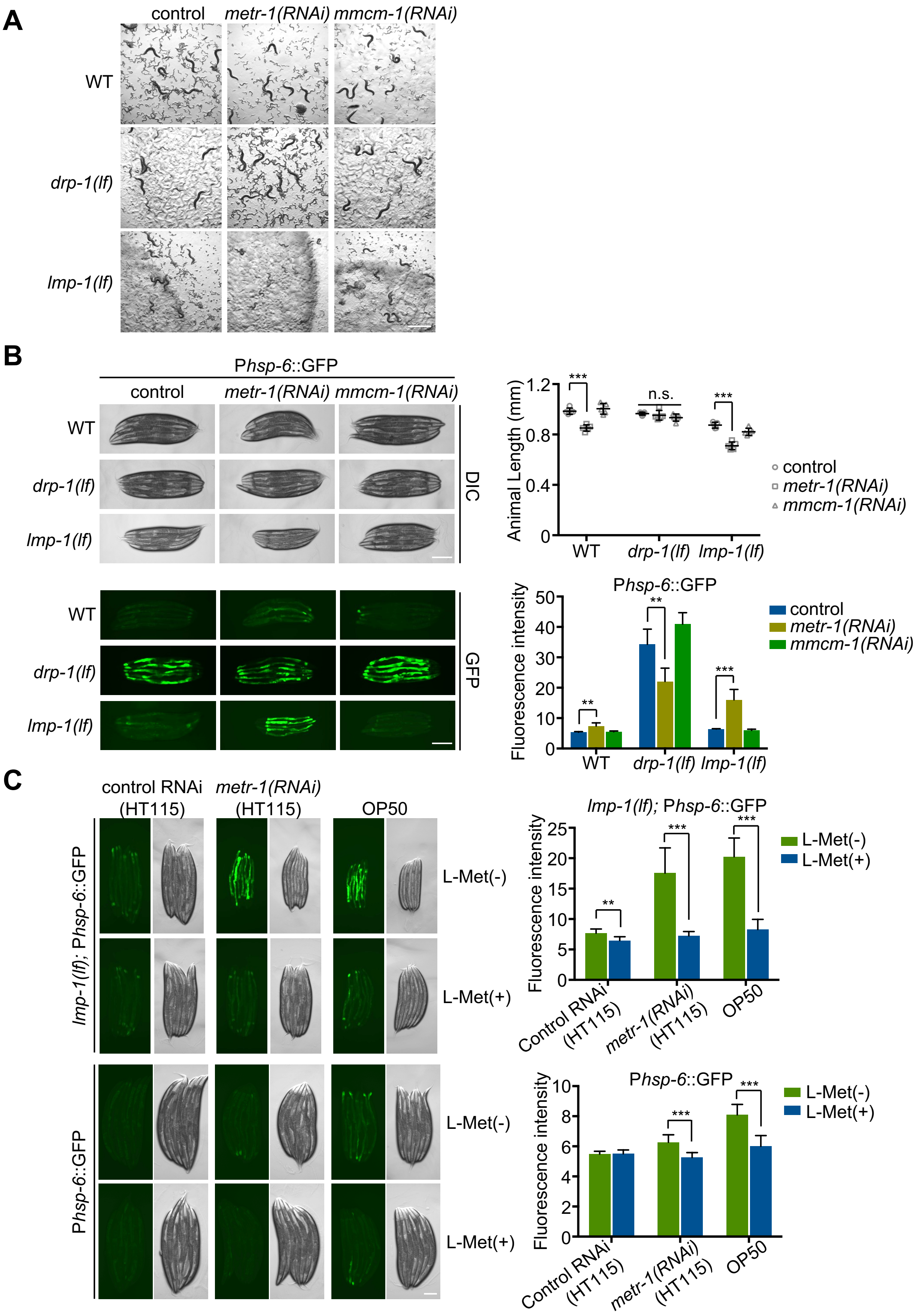

### Supplementary Figure 12

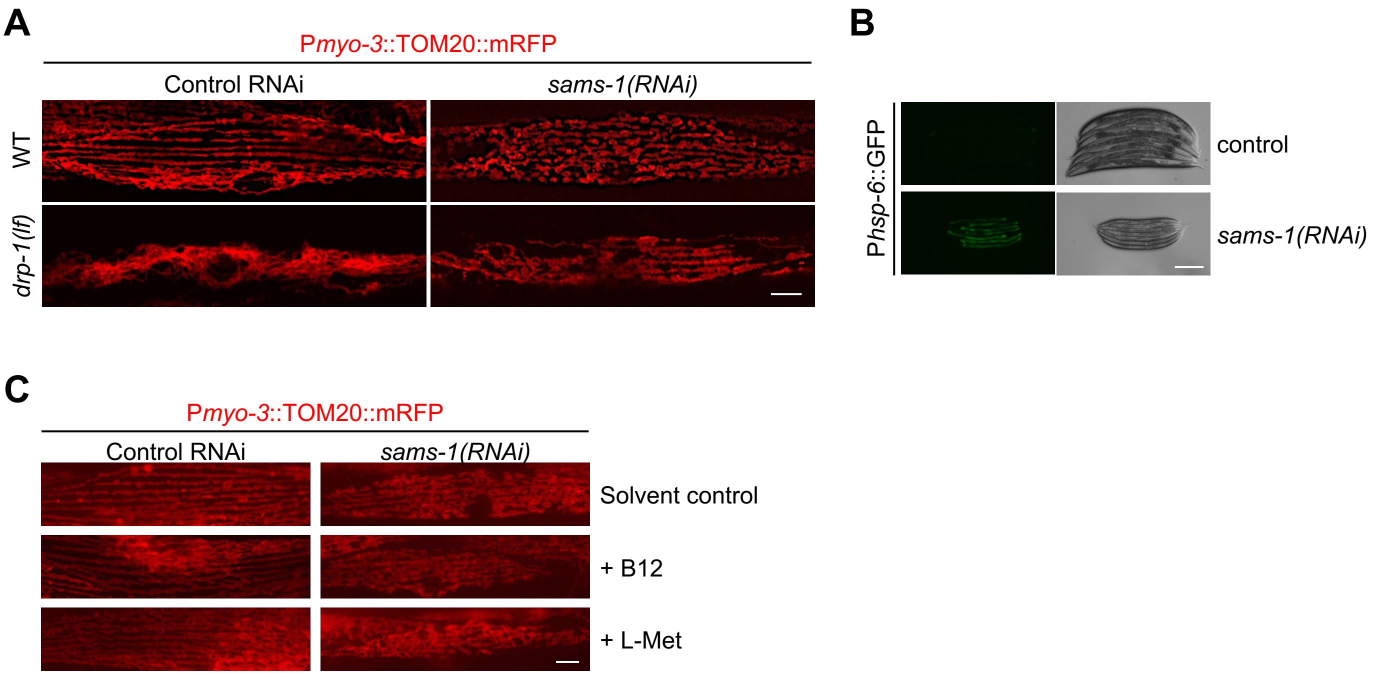

#### Supplementary Figure 13

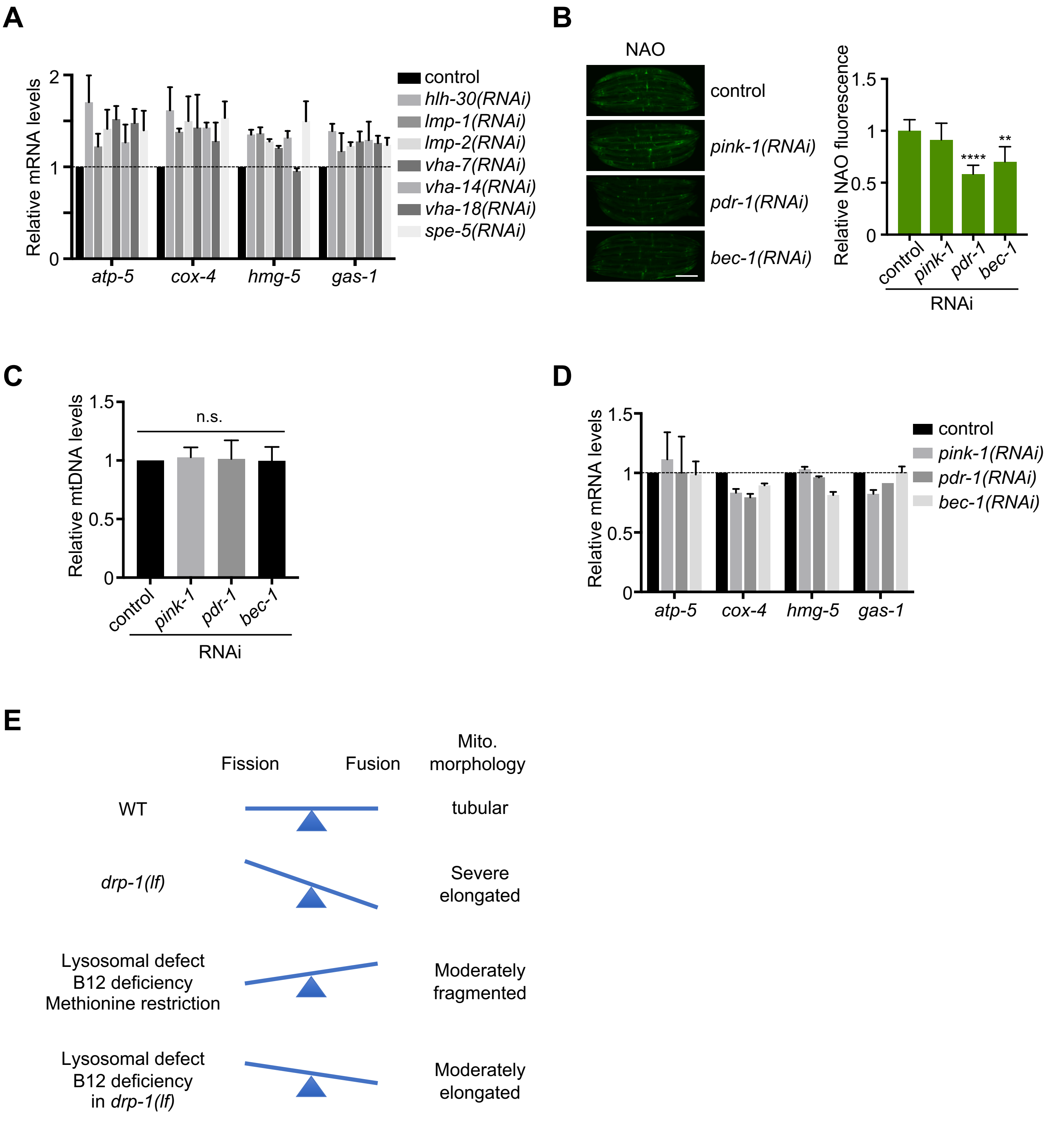
